# Molecular basis for plasma membrane recruitment of PI4KA by EFR3

**DOI:** 10.1101/2024.04.30.587787

**Authors:** Sushant Suresh, Alexandria L Shaw, Joshua G Pemberton, Mackenzie K Scott, Noah J Harris, Matthew AH Parson, Meredith L Jenkins, Pooja Rohilla, Alejandro Alvarez-Prats, Tamas Balla, Calvin K Yip, John E Burke

## Abstract

The lipid kinase phosphatidylinositol 4 kinase III alpha (PI4KIIIα/PI4KA) is a master regulator of the lipid composition and asymmetry of the plasma membrane. PI4KA exists primarily in a heterotrimeric complex with its regulatory proteins TTC7 and FAM126. Fundamental to PI4KA activity is its targeted recruitment to the plasma membrane by the lipidated proteins EFR3A and EFR3B. Here, we report a cryo-EM structure of the C-terminus of EFR3A bound to the PI4KA-TTC7B-FAM126A complex, with extensive validation using both hydrogen deuterium exchange mass spectrometry (HDX-MS), and mutational analysis. The EFR3A C-terminus undergoes a disorder-order transition upon binding to the PI4KA complex, with an unexpected direct interaction with both TTC7B and FAM126A. Complex disrupting mutations in TTC7B, FAM126A, and EFR3 decrease PI4KA recruitment to the plasma membrane. Multiple post-translational modifications and disease linked mutations map to this site, providing insight into how PI4KA membrane recruitment can be regulated and disrupted in human disease.

**One sentence summary:** The cryo-EM structure of the C-terminus of EFR3A bound to the PI4KA complex reveals the molecular mechanism underlying PI4KA membrane localisation, and provides novel insight into PI4KA regulation and its involvement in human disease.

## Introduction

Phosphoinositide lipids play numerous roles in myriad cellular functions (*1–3*). One of the most abundant phosphoinositide lipids is the molecule phosphatidylinositol 4-phosphate (PI4P) (*4*), which can be generated by four different lipid kinases in humans (encoded by the genes *PI4KA, PI4KB, PI4K2A,* and *PI4K2B*), with PI4KA generating the primary pool of PI4P at the plasma membrane (PM) (*5*). PI4P is essential for defining the identity of the PM, as it is the precursor for the signaling lipids phosphatidylinositol (4,5)-bisphosphate (PI(4,5)P_2_) and phosphatidylinositol (3,4,5)-trisphosphate (PIP_3_), regulates the non-vesicular transport of phosphatidylserine (*6*), and is also a substrate of the phospholipase C (PLC) pathway (*7*). Dysregulation of PI4KA is involved in multiple human diseases (*8*), with loss of function mutations in PI4KA or its regulatory proteins being causative of neurological (*9*, *10*), immunological (*11*), and gastrointestinal disorders (*12*, *13*). PI4KA and its regulatory proteins have also been shown to be critical for cancer growth (*14*, *15*) and viral infection (*16*).

PI4KA shares an evolutionary history with the class I-III PI 3-kinases (PI3Ks) and PI4KB, with a central feature of regulation for all of these enzymes being that they are recruited to membrane surfaces by specific lipid or protein binding partners (*17*). PI4KA primarily exists in cells as a complex with the regulatory proteins TTC7 (two isoforms TTC7A and TTC7B) and FAM126 (two isoforms FAM126A and FAM126B) (*18*, *19*). PI4KA and TTC7 are conserved throughout all eukaryotes, while FAM126 is found in most metazoans, with it not being found in fungi, but found in organisms containing a simple nervous system (*20*). Analysis of the PI4KA-TTC7-FAM126 complex using cryogenic electron microscopy (cryo-EM) and hydrogen deuterium exchange mass spectrometry (HDX-MS) showed that it assembles into a large ∼750 kDa complex (*18*, *21*). PI4KA self-assembles through its dimerization domain leading to a dimer of heterotrimers, which we will refer to as the PI4KA complex. FAM126 does not bind to PI4KA directly but instead acts as a stabilizer of TTC7, which binds to PI4KA (*19*). The most well-established mechanism of membrane recruitment is through the binding of the palmitoylated protein EFR3 to TTC7 (*5*). However, the exact molecular mechanism underlying EFR3-mediated PM recruitment of the PI4KA complex has remained elusive (*5*, *22*, *23*).

The interaction between EFR3 and TTC7 has been extensively studied in yeast, where the protein Ypp1 (yeast ortholog of TTC7) binds to the disordered C-terminus of Efr3 (yeast homolog of EFR3) (*24*, *25*). In yeast, this interaction is regulated by phosphorylation of the C-terminus of Efr3, with phosphorylation disrupting Ypp1-mediated PM recruitment of Stt4 (yeast homolog of PI4KA) (*25*). It has been challenging to extrapolate these results to derived eukaryotic lineages, including humans, as the C-terminus of EFR3 is only partially conserved between yeast and humans. As PI4KA activity needs to be upregulated upon PLC signaling to maintain the PM pool of PI4P (*26*), there likely are unknown regulatory mechanisms controlling EFR3-mediated recruitment in vertebrates. Defining the molecular mechanism of EFR3-mediated PM recruitment of PI4KA is essential to understanding possible regulatory mechanisms controlling the activity of the PI4KA complex through association with the PM.

To fully explore the molecular mechanisms of how EFR3 can bind to and regulate the PI4KA complex, we have used a synergistic application of cryo-EM and HDX-MS. Our structural analyses reveal that EFR3A interacts with the PI4KA complex through an evolutionarily conserved region within the C-terminus of EFR3A. Unexpectedly, the C-terminus of EFR3A forms an extended helical interface toward both TTC7B and FAM126A. This interface was validated by both HDX-MS and mutational analysis. The development of a live-cell bioluminescence resonance-energy transfer (BRET)-based assay of PI4KA recruitment to the PM as well as confocal imaging show the key functional role of the EFR3 C-terminus in membrane recruitment of the PI4KA complex. Overall, our findings provide useful insight into the molecular and structural basis of PI4KA regulation at the PM.

## Results

### Cryo-EM analysis of the EFR3A-PI4KA complex

Investigation into the molecular basis of how EFR3A regulates PI4KA required the production of a stable PI4KA-TTC7B-FAM126A complex. This was achieved by co-expression of full-length PI4KA and TTC7B, together with a truncated FAM126A lacking the disordered C-terminus (1-308, FAM126A ΔC) (**Fig. 1A**). This complex has been structurally characterized (*18*), is homogenous, and elutes off gel filtration consistent with the formation of a dimer of heterotrimers. For simplicity, this complex will be referred to as the PI4KA complex throughout the manuscript. It has previously been established that in yeast the disordered C-terminus of EFR3 mediates recruitment of the TTC7 ortholog Ypp1 (*25*). To identify the related site in human EFR3A we carried out a bioinformatic analysis of sequence conservation. While there were no regions that were strictly conserved between yeast and humans (**Supplemental Fig 1)**, there was a region that was highly conserved among chordates in the C-terminus. We generated multiple constructs spanning this region for expression in *E. coli*, with the most stable and homogenous being a construct spanning residues 721-791 of EFR3A (**Fig. 1A**).

**Figure 1.**
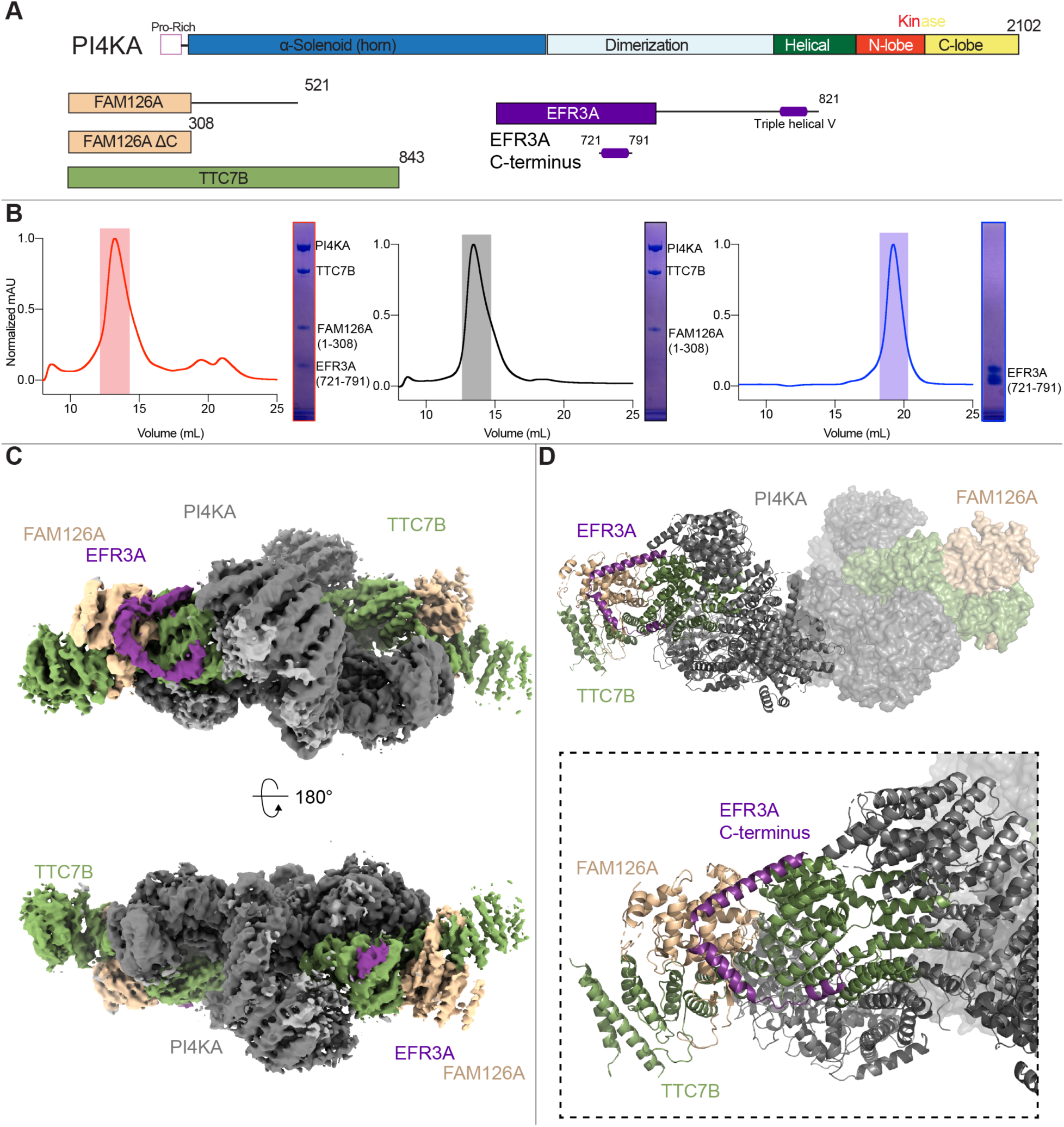
Cryo-EM analysis of EFR3 binding to the PI4KA complex. **A.** Domain schematics of the full-length PI4KA complex and EFR3A constructs used in this paper. Constructs used in this paper are PI4KA/TTC7B/FAM126A ΔC (referred to as PI4KA complex) and EFR3A (721-791) referred to as EFR3A. **B.** Size exclusion chromatography traces of (L) PI4KA complex/EFR3A, (M) PI4KA complex, and (R) EFR3A, with corresponding SDS-PAGE gels to show protein present in the highlighted size exclusion chromatography (SEC) peaks. **C.** Cryo-EM density map of the PI4KA complex bound to EFR3A. EFR3A was bound to both sides of the hetero-trimer, however, the full triple helical V was only visible in one of the two dimers of hetero-tetramers (see left side of the molecule in the top panel). **D.** Molecular model of the triple helical V in the C-terminus of EFR3A (724-787) bound to the PI4KA complex, with a zoom-in highlighting EFR3A contacts with both TTC7B and FAM126A.

To validate that this EFR3A fragment could interact with the PI4KA complex, size exclusion chromatography was used to determine whether complex formation would occur. When co-incubated together, the PI4KA complex co-eluted with the EFR3A C-terminal fragment, with elution volume and SDS-PAGE analysis consistent with a dimer of hetero-tetramers (**Fig. 1B**). We then carried out cryo-EM analysis of this reconstituted complex. While the protein specimen appeared homogeneous on negative stain EM grids, vitrification of this protein product yielded aggregated particles at the air-water interface. To address this problem, we subjected the complex to limited BS^3^ (bis(sulfosuccinimidyl)suberate) chemical crosslinking followed by size exclusion chromatography, with this complex eluting at a volume similar to the non-crosslinked complex.

Using this sample, we were able to generate a cryo-EM reconstruction of the PI4KA complex bound to the C-terminus of EFR3A at a nominal resolution of 3.48Å from 67,563 particles (**Fig. 1C, Supplemental Fig 2**). Refinement yielded large differences in local resolution between the two hetero-tetramers. We will therefore focus our descriptions on the better-resolved hetero-tetramer within the assembly. (**Supplemental Fig 2C**). Unexpectedly, our structure showed that the EFR3A C-terminus binds to both TTC7B and FAM126A, forming a V-shape composed of three alpha helices. The local resolution of the first interfacial α-helix in EFR3A spanning residues 728-735 was sufficient for unambiguous manual model building into density (**Supplemental Fig 3**). The local resolution of the 2nd and 3rd helices of EFR3A was insufficient for reliable manual model building, so we used rigid body refinement of an AlphaFold3 prediction (*27*, *28*) to model these helices (**Supplemental Fig 3**). Overall, we were able to build a molecular model of residues 724-787 of EFR3A bound to TTC7B-FAM126A (**Fig. 1D**). No direct interactions were seen between the EFR3A C-terminus and PI4KA, with no major conformational changes in the PI4KA subunit. The interface between EFR3A and FAM126A was unexpected, as both EFR3 and TTC7 are conserved from yeast to humans, while FAM126 is conserved only in metazoans (*20*).

### HDX-MS analysis of the EFR3A-PI4KA complex

To further understand the dynamics of the interaction between the PI4KA complex and EFR3A, we employed HDX-MS, a powerful technique to investigate protein conformational dynamics (*29–31*). HDX-MS experiments were performed on the PI4KA complex with and without EFR3A. Deuterium incorporation was measured over a range of time points (30s, 300s, 3000s, 10000s) and the mass shift upon deuterium incorporation were analysed via mass spectrometry, with significant differences in exchange being defined as changes greater than >0.45 Da and >5% at any time point with unpaired t-test values of p<0.01.

Multiple regions in both TTC7B and FAM126A showed significant decreases in exchange upon binding to EFR3A. These regions included residues 89-102 in FAM126A and residues 539-544 in TTC7B (**Fig. 2A+B, Supplemental Fig 4**), which were all localized at the interface observed by cryo-EM. No significant changes in exchange were observed in PI4KA, consistent with the lack of direct contacts observed in the cryo-EM structure.

**Figure 2:**
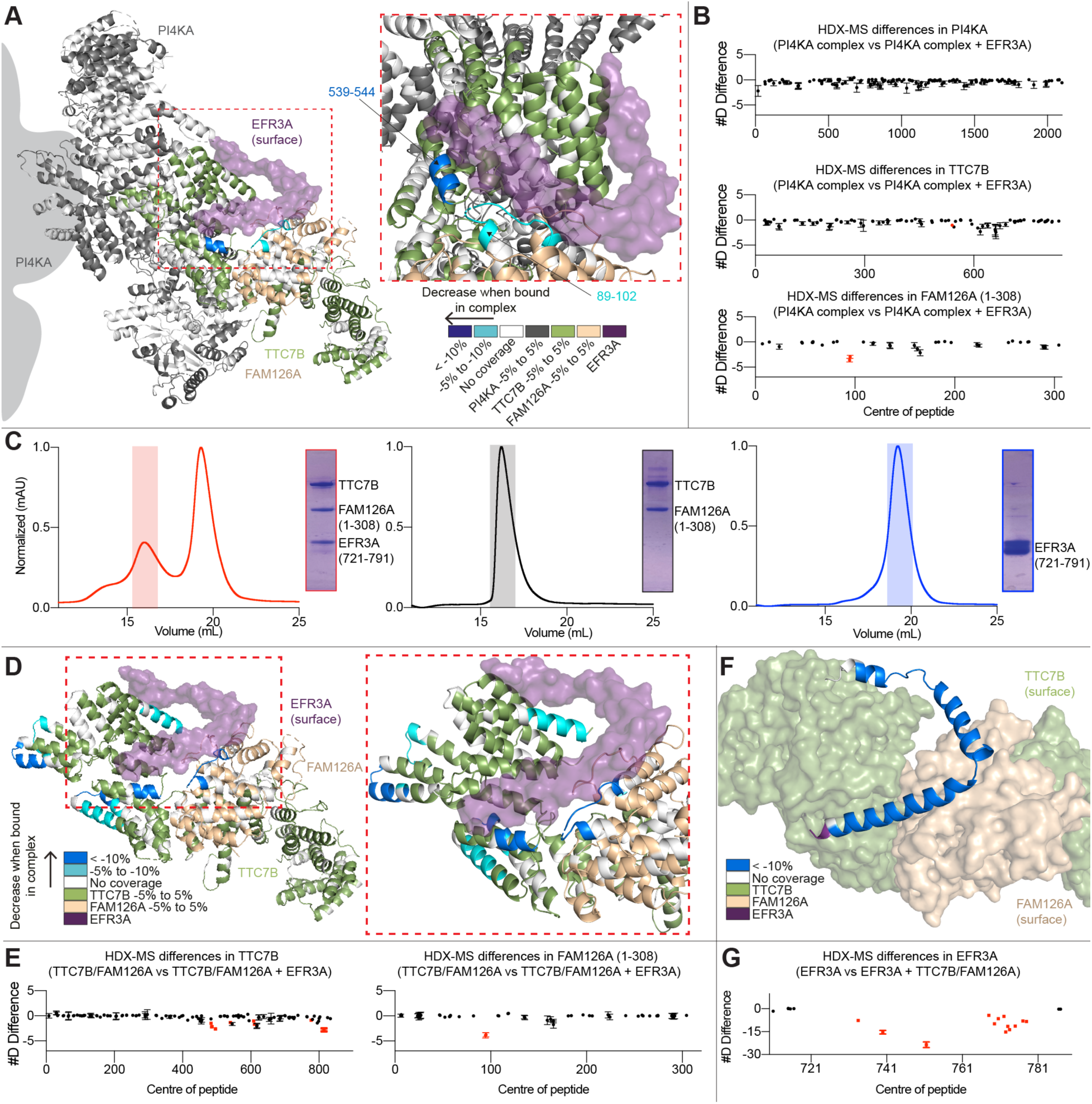
HDX-MS analysis of the interaction of EFR3A with TTC7B and FAM126A. **A.** Residues showing significant differences in deuterium exchange (defined as >5% 0.45 Da, and p<0.01 in an unpaired two-tailed t-test at any time point) upon PI4KA complex binding to EFR3A. Differences are mapped on a structural model of the high local resolution half, with a zoom-in on the interface. Differences are indicated by the legend. **B.** Sum of the number of deuteron differences of PI4KA complex upon formation with EFR3A analysed over the entire deuterium exchange time course for the PI4KA complex. Each point is representative of the centre residue of an individual peptide. Peptides that met the significance criteria described in A are coloured red. Error is shown as the sum of standard deviations across all 4 time points (SD) (n=3 for each condition in each timepoint). **C.** Size exclusion chromatography traces of (L) TTC7B-FAM126A ΔC -EFR3A, (M) TTC7B-FAM126A ΔC, and (R) EFR3A, with corresponding SDS-PAGE gels to show protein present in the highlighted SEC peaks. **D.** TTC7B-FAM126A residues showing significant differences in deuterium exchange (defined as >5% 0.45 Da, and p<0.01 in an unpaired two-tailed t-test at any time point) upon formation of the TTC7B-FAM126A-EFR3A complex. Differences are mapped on a structural model of the high local resolution half removing PI4KA, with a zoom in on the interface. Differences are indicated by the legend. **E.** Sum of the number of deuteron differences of TTC7B and FAM126A upon complex formation with EFR3A analysed over the entire deuterium exchange time course for the dimer. Each point is representative of the centre residue of an individual peptide. Peptides that met the significance criteria described in D are coloured red. Error is shown as the sum of standard deviations across all 5 time points (SD) (n=3 for each condition in each timepoint). **F.** EFR3A residues showing significant differences in deuterium exchange (defined as >5% 0.45 Da, and p<0.01 in an unpaired two-tailed t-test at any time point) upon formation of the TTC7B-FAM126A-EFR3A complex. Differences are mapped on a structural model of the high local resolution half removing PI4KA zooming in on the interface. Differences are indicated by the legend. The construct used has overlapping MBP peptides that do not contain EFR3A residues, therefore peptides that start before 721 are not EFR3A sequence. **G.** Sum of the number of deuteron differences of EFR3A upon complex formation with TTC7B-FAM126A analysed over the entire deuterium exchange time course for the EFR3A. Each point is representative of the centre residue of an individual peptide. Peptides that met the significance criteria described in F are coloured red. Error is shown as the sum of standard deviations across all 4 time points (SD) (n=3 for each condition in each timepoint). Individual deuterium exchange curves for significant differences for all conditions are shown in Supplemental Fig 4.

Together, the cryo-EM and HDX-MS data suggested that EFR3A would bind similarly to an isolated dimer of TTC7B-FAM126A ΔC. We purified TTC7B-FAM126A ΔC, which will be referred to as TTC7B-FAM126A, from *E. coli*, with gel filtration profiles in the presence of the C-terminus of EFR3A consistent with the formation of a stable trimer (**Fig. 2C**). To investigate the implications of the lack of PI4KA on conformational dynamics, we used HDX-MS to compare TTC7B-FAM126A in the presence and absence of the C-terminus of EFR3A. HDX-MS experiments were measured over 5 time points (3s, 30s, 300s, 3000s and 10,000s at 18°C). We observed significant decreases in HDX in the same sites on TTC7B and FAM126A as observed in the PI4KA complex, with additional regions spanning the entire EFR3A interface, as well as some allosteric differences distant from the EFR3A binding site (**Fig. 2D-E, Supplemental Fig 4**). To map changes in deuterium incorporation in EFR3A, we used a different EFR3A C-terminus construct containing residues 721-791 of EFR3A with an N-terminal maltose binding protein (MBP) tag, with HDX-MS experiments performed on MBP-EFR3A apo and in complex with TTC7B-FAM126A. Deuterium incorporation was measured over 4 time points (3s, 30s, 300s at 18°C and 3s at 0°C). Peptides spanning all three helices in EFR3A showed a significant decrease in HDX, further validating the interface observed in the cryo-EM structure (**Fig. 2F-G, Supplemental Fig 4**). We carried out biolayer interferometry (BLI) measurements of TTC7B-FAM126A binding to the C-terminus of EFR3A, with this interface having a *K*_D_ of ∼63 ± 13 nM (**Fig. 3A**).

**Figure 3:**
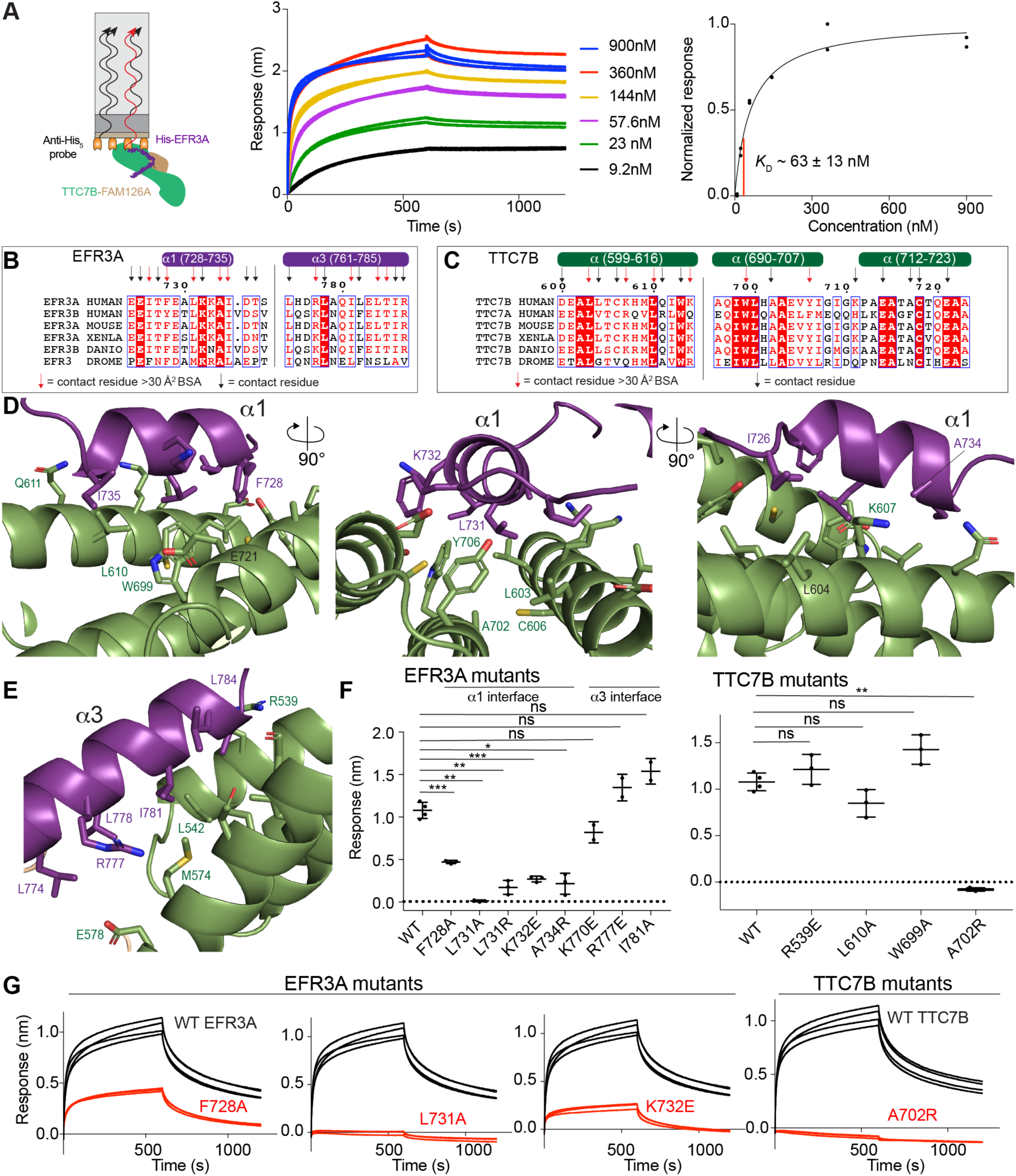
Molecular basis of EFR3A binding to TTC7B. **A.** Cartoon schematic (L) of a bio-layer interferometry (BLI) experiment showing binding of immobilized His-EFR3A (721-791) to TTC7B-FAM126A. Association and dissociation curves (M) for the binding of His-EFR3A (721-791) to different concentrations of TTC7B-FAM126A (10-2500 nM). The experiment was carried out in duplicate, with all data shown. Normalized BLI response versus concentration of TTC7B-FAM126A (R), with *K*_D_ estimated by one site specific non-linear regression. Each data point is shown (n=2). **B.** Multiple sequence alignment of EFR3A from *Homo sapiens, Mus musculus, Xenopus laevis, Danio rerio, Drosophila melanogaster*. EFR3A secondary structure of α1 and α3 helices are annotated above the alignment. Contact residues within 6 Å of TTC7B are annotated using arrows. Contact residues with BSA > 30 Å^2^ are annotated using red arrows. **C.** Multiple sequence alignment of TTC7B from *Homo sapiens, Mus musculus, Xenopus laevis, Danio rerio, Drosophila melanogaster*. TTC7B secondary structure is annotated above the alignment. Contact residues within 6 Å of EFR3A are annotated using arrows. Contact residues with BSA > 30 Å^2^ are annotated using red arrows. **D.** Zoomed in cartoon view of the EFR3A α1-TTC7B interface with putative interfacial residues labelled. EFR3A and TTC7B are coloured according to in-figure text. **E.** Zoomed in cartoon view of the EFR3A α3-TTC7B interface with putative interfacial residues labelled. EFR3A and TTC7B are coloured according to in-figure text. **F.** Maximum BLI response of various EFR3A and TTC7B mutants compared to WT. Error is shown as standard deviation (n=3) with two-tailed p values indicated as follows: **p<0.001; ***p<0.0001; not significant (ns) > 0.01. **G.** Raw BLI association and dissociation curves of EFR3A and TTC7B mutants compared to WT. His-EFR3A was loaded on the anti-penta-His tip at 200 nM and dipped in TTC7B-FAM126A at 500 nM. Raw BLI curves of all mutants in Supplemental Fig 5.

### Molecular details of the EFR3A interface with TTC7B

Our cryo-EM structure revealed that EFR3A engages TTC7B and FAM126A in an extended interface composed of three α-helices from EFR3A oriented in a V, resulting in a buried surface area of ∼1500 Å^2^. This interface is primarily hydrophobic, with the interface between EFR3A and TTC7B being ∼800 Å^2^ and consists of the first EFR3A α-helix (residues 728-735) and the latter half of the 3^rd^ EFR3A α-helix (761-785) (**Fig. 3B-C**). The first α-helix is likely the most stable as this is the only region of EFR3A visible in both halves of the cryo-EM map and is also the region with the highest local resolution in the electron density (**Fig 2C)**. The α1-helix in EFR3A is amphipathic, with the hydrophobic face primarily interacting with hydrophobic residues in three helices in TTC7B (residues 599-616, 690-707, and 712-723). The interacting residues in the α1 and α3 helices of EFR3A are strongly conserved throughout evolution in vertebrates, with partial conservation in metazoans, and very limited conservation in yeast (**Fig 3B and Supplemental Fig 1**). There are some notable differences in interfacial residues between EFR3A and EFR3B that may alter binding affinity, potentially highlighting isoform-specific regulatory differences.

We generated several point mutations on both EFR3A and TTC7B and used Bio-Layer Interferometry (BLI) to determine their effect on complex formation (**Fig 3F-G and Supplemental Fig 5**). Mutation of residues in the α1 helix in EFR3A, including F728A, L731A, K732E, and A734R led to decreased binding, with the L731A mutation causing the largest disruption. Mutation of a corresponding residue in TTC7B (A702R) also led to an almost complete disruption of EFR3A binding, with removal of hydrophobic residues showing a minimal effect on binding affinity. Mutations of residues in the α3 helix of EFR3A, or corresponding residues in TTC7B showed no significant effect on binding, highlighting that the α1 helix of EFR3A is indispensable for binding, with a limited role of the C-terminal half of the α3 helix in EFR3A.

### Molecular details of the EFR3A interface with FAM126A

It was unexpected that there would be an extensive interface between EFR3A and FAM126A, as FAM126A has not been previously annotated as a critical partner in EFR3-mediated PM recruitment of PI4KA. There is an extended evolutionarily conserved interface between EFR3A and FAM126A (**Fig. 4A-C**). This interface in EFR3A is composed of the C-terminal region of the α2-helix, the N-terminal region of the α3-helix, and the residues between these two helices. This packs up against two α-helices in FAM126A (53-66 and 97-117).

**Figure 4.**
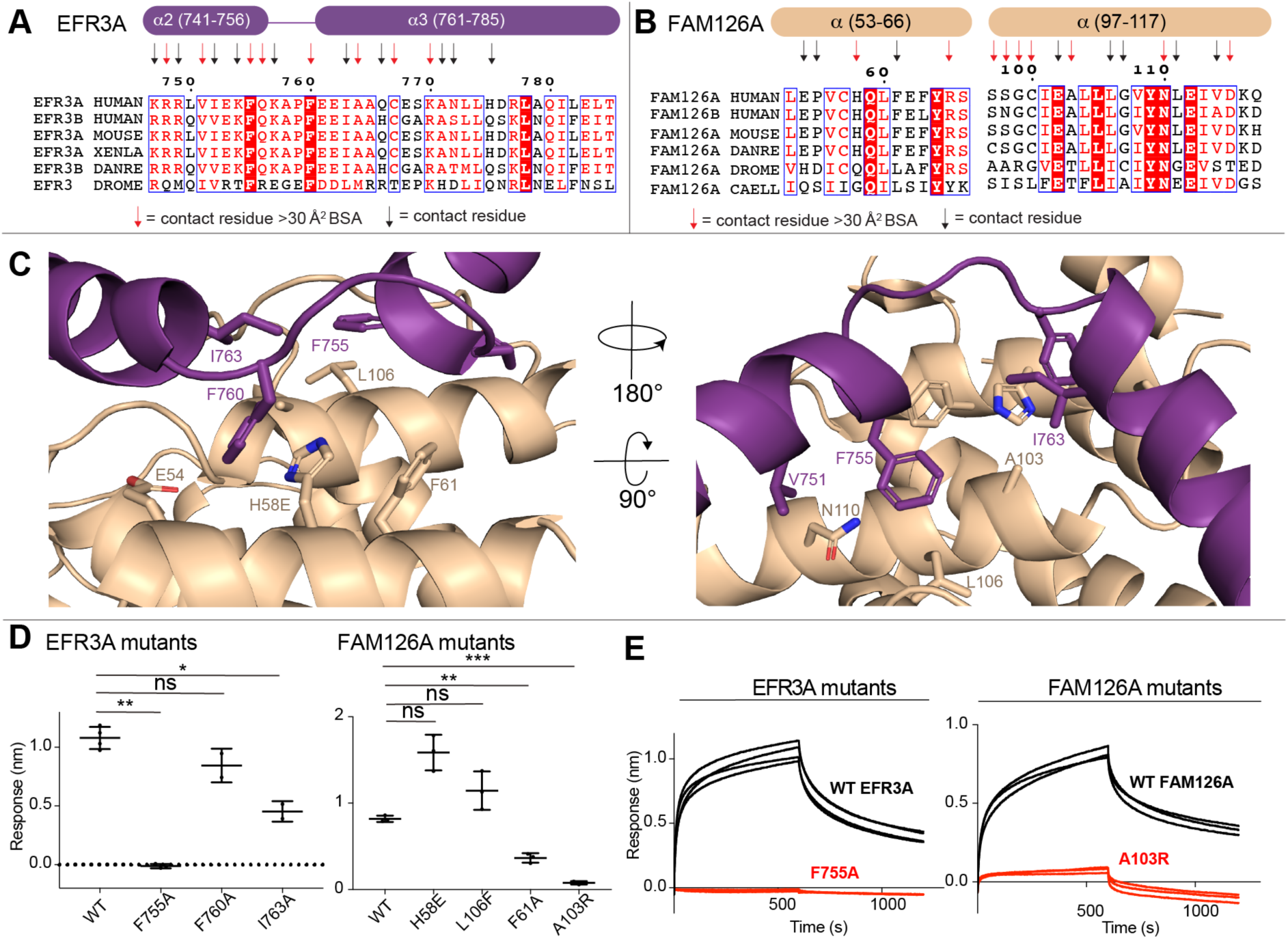
Mutational analysis validates the EFR3A-FAM126A binding interface. **A.** Multiple sequence alignment of EFR3A from *Homo sapiens, Mus musculus, Xenopus laevis, Danio rerio and Drosophila melanogaster*. EFR3A secondary structure of α2 and α3 helices are annotated above the alignment. Contact residues within 5 Å of FAM126A are annotated using arrows. Contact residues with BSA > 30 Å^2^ are annotated using red arrows. **B.** Multiple sequence alignment of FAM126A from *Homo sapiens, Mus musculus, Danio rerio, Drosophila melanogaster and Caenorhabditis elegans*. FAM126A secondary structure is annotated above the alignment. Contact residues within 5 Å of EFR3A are annotated using arrows. Contact residues with BSA > 30 Å^2^ are annotated using red arrows. **C.** Zoomed in cartoon view of the EFR3A-FAM126A interface with putative interfacial residues labelled. EFR3A and FAM126A are coloured according to in-figure text. **D.** Maximum BLI response of various EFR3A and FAM126A mutants compared to WT. Error is shown as standard deviation (n=3) with p values indicated as follows: *p<0.01; **p<0.001; ***p<0.0001; not significant (ns) > 0.01. **E.** Raw BLI association and dissociation curves of EFR3A and FAM126A mutants compared to WT. His-EFR3A was loaded on the anti-penta-His tip at 200 nM and dipped in TTC7B-FAM126A at 500 nM. Raw BLI curves of all mutants in Supplemental Fig 5.

We generated several point mutations at the EFR3A-FAM126A interface and used BLI to determine their effect on complex formation (**Fig 4D-E and Supplemental Fig 5**). Mutation of residues F755A and I763A localized at the interface of helix α2 and α3 led to decreased binding, with the F755A mutation causing the largest disruption. Mutation of corresponding residues in FAM126A (F61A and A103R) also led to significant disruption of EFR3A binding. This validates the previously unidentified critical role of FAM126A in EFR3A binding, with this likely having important implications for PI4KA regulation.

### BRET analysis of EFR3-mediated plasma membrane recruitment

To define the functional role of the identified interface between EFR3 and TTC7-FAM126 in mediating PM localization of PI4KA, we developed a unique bioluminescence resonance energy transfer (BRET) assay. These quantitative measurements used a single-plasmid design containing a PM (Lyn Kinase N-Terminus; L10)-anchored BRET acceptor (mVenus) and a nano-Luciferase (nLuc)-tagged PI4KA, which were separated by a self-cleaving tandem viral 2A peptide sequence (tPT2A) to produce each protein in transfected cells at a fixed stoichiometry (**Fig. 5A**). Upon addition of a cell-permeable luciferase substrate (coelenterazine h; final concentration 5 μM), the signal from the nLuc-generated luminescence and resulting mVenus fluorescence can be measured in entire populations of live cells using a multimode microplate reader. Experiments were carried out by transfecting this PM-PI4KA^BRET^ biosensor (L10-mVenus-tPT2A-nLuc-PI4KA) along with a single-plasmid encoding full length epitope-tagged EFR3B, TTC7B, and FAM126A which also contains unique P2A and T2A splitting sequences between the specific components (EFR3B-P2A-TTC7B-T2A-FAM126A), as we have previously described

**Figure 5.**
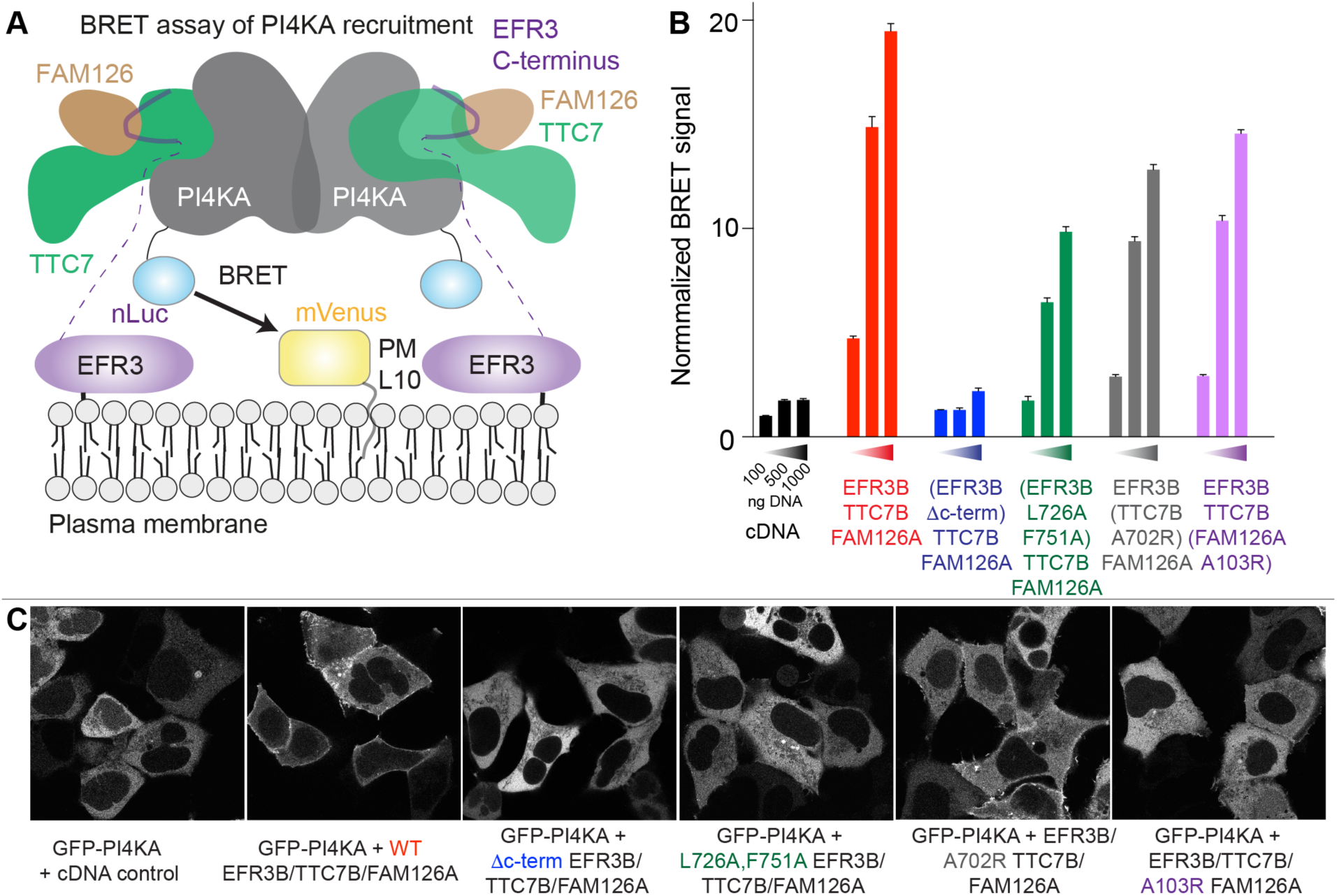
The C-terminus of EFR3A mediates plasma membrane recruitment of PI4KA. **A.** Cartoon depicting the quantitative BRET-based PI4KA recruitment assay. Briefly, the PM-anchored BRET acceptor (L10-mVenus) will only be near the nLuc-tagged PI4KA BRET-donor, and thereby efficiently increase the relative BRET signal, if co-assembled with the requisite molecular partners for PM targeting. **B.** Normalized BRET signal measured from HEK293A cell populations (∼0.75 x 10^5^ cells/well) expressing a fixed amount of the PM-PI4KA^BRET^ biosensor (L10-mVenus-tPT2A-NLuc-PI4KA; 500 ng/ well) together with increasing amounts (100, 500, or 1000 ng/well) of constructs consisting of either an empty vector (black), wild-type control (red; EFR3B-P2A-TTC7B-T2A-FAM126A), or the indicated mutants of the PI4KA complex components (blue, EFR3B Δc-term(1-716)-P2A-TTC7B-T2A-FAM126A; green, EFR3B^L726A,F751A^-P2A-TTC7B-T2A-FAM126A; grey, EFR3B-P2A-TTC7B^A702R^-T2A-FAM126A; and, magenta, EFR3B-P2A-TTC7B-T2A-FAM126A^A103R^). For all treatments, BRET values were normalized relative to an internal basal BRET control (PM-PI4KA^BRET^ biosensor expressed without the empty vector or EFR3B-P2A-TTC7B-T2A-FAM126A plasmid added) and are presented as the summary of triplicate wells measured for each treatment condition in three independent biological replicates (n = 3, with normalized signals from 9 total wells averaged). Normalized BRET signals were then expressed and graphed as fold-increases relative to the 100 ng/well pcDNA3.1 control. **C.** Representative confocal images of live HEK293A cells co-expressing a fixed amount (250 ng) of EGFP-PI4KA (greyscale) together with the indicated constructs (500 ng/each). The sequence and colour-coding of the experimental treatments are matched with the BRET measurements in (**B**).

(*32*).

Live-cell experiments were carried out using HEK293A cells expressing a fixed amount of the PM-PI4KA^BRET^ biosensor (500ng/well), along with increasing amounts (100, 500, and 1000 ng/well) of the EFR3B-TTC7B-FAM126A plasmid. Importantly, we saw a clear dose-dependent increase in the basal BRET signal when full-length EFR3B, TTC7B and FAM126A were co-expressed (red bars), consistent with an increase to the PM recruitment of PI4KA (**Fig. 5B**). We performed a control experiment where the C-terminus of EFR3B was truncated (EFR3B Δc-term, where residues 716-817 were truncated) and saw no significant increase in the BRET signal (blue bars) over an empty-vector control (black bars). This is consistent with the C-terminus of EFR3 being required for EFR3 mediated PM recruitment of PI4KA. Subsequently, we incorporated complex disrupting mutations in EFR3B (L726A, F751A; green bars), TTC7B (A702R; grey bars), or FAM126A (A103R; magenta bars), and saw a significantly reduced BRET signal across the various levels of relative plasmid expression when compared to the wild-type controls. However, all these mutants still had a measurable BRET signal above that of the truncated EFR3B control (**Fig. 5B**). This indicates the importance of avidity in PI4KA recruitment, as none of these mutations affecting the complex interface were able to completely abrogate PI4KA localization to the PM. In parallel with this BRET-based approach, we also used confocal microscopy for live-cell imaging studies to examine the subcellular localization of an EGFP-tagged PI4KA construct after co-expression of EFR3B, TTC7B, and FAM126A. Similar to our quantitative BRET-based measurements, we only saw robust PM localization of EGFP-PI4KA when co-expressed together with the wild-type EFR3B, TTC7B and FAM126A, and very limited PM localization of EGFP-PI4KA in conjunction with the mutant or truncated versions of EFR3B, TTC7B, and FAM126A (**Fig. 5C**).

## Discussion

Understanding how PI4KA is recruited, activated, and regulated at the PM is essential in developing therapeutic strategies for targeting human diseases associated with loss-of-function of PI4KA. PI4KA, TTC7A and FAM126A mutations have been identified in immunological, neurological, gastrointestinal, and developmental disorders (*9–12*, *19*, *33*). Some cancers also require EFR3 and PI4KA activity for their aggressive growth (*14*, *15*). Here, we report the architecture of the EFR3A complex that is formed with PI4KA-TTC7B-FAM126A, revealing novel insights into the molecular mechanisms underlying PM recruitment of PI4KA.

Our cryo-EM and HDX-MS analyses reveal that EFR3A forms an extended interface with both the TTC7 and FAM126 regulatory proteins. Previous analysis of evolutionarily conserved surfaces in TTC7 and FAM126 had led to the proposal of an EFR3 interface composed of both proteins (*18*), and our work provides the first experimental evidence of the cooperation between TTC7 and FAM126 for EFR3 binding. This was validated by complex-disrupting mutations in EFR3A at both the FAM126A and TTC7B interfaces, as well as mutations in FAM126A and TTC7B at the interface with EFR3A. There was no difference in the conformation of PI4KA upon binding EFR3A in solution, suggesting that there are no allosteric conformational changes upon EFR3A binding. However, further biochemical and biophysical analysis of membrane resident full-length EFR3 recruiting the PI4KA complex to membranes will be required to fully understand the conformational dynamics of PI4KA upon membrane recruitment.

An important implication of this structure is that the two EFR3 isoforms may be able to be selectively recruited to specific combinations of TTC7 and FAM126 isoforms. While the binding interfaces are conserved in different isoforms, this conservation is not absolute, allowing for the possibility of unique binding affinities. TTC7A has a 50% sequence similarity with TTC7B, while FAM126A has a 54% sequence similarity to FAM126B. Some of the most conserved regions are located at the EFR3 interface. For EFR3, the C-terminus has a 67% sequence similarity between EFR3A and EFR3B, with most of the interfacial residues being strongly conserved. It is still not known if these sequence differences drive different roles of TTC7/FAM126/EFR3 isoforms or if these differences are due to variable tissue expression. Isoforms of TTC7, FAM126 and EFR3 are differentially expressed in diverse tissues, with many tissues expressing multiple isoforms. The isoform-specific roles of the PI4KA regulatory proteins are still not fully elucidated.

The potential for unique molecular roles for the isoform-specific PI4KA complexes has been recently highlighted for TTC7A and TTC7B, with patients having disease-linked loss of function mutations in TTC7A in tissues that express both TTC7A and TTC7B, where it appears that TTC7B cannot compensate for the loss of TTC7A. This may be driven by differences in TTC7A/TTC7B expression during embryogenesis, where TTC7A is widely expressed, and TTC7B is primarily expressed in brain tissues (*34*). As the experimental structure of TTC7A is unknown, we examined AlphaFold3 predicted models of TTC7A compared to our structure, with a focus on potential differences at the EFR3A interface. Intriguingly, there is a weakly predicted TTC7A helix (residues 678-690) in the same location as the α1 helix of EFR3A, with this region being only weakly conserved in TTC7B (**Supplemental Fig 6A**) (*35*). In both TTC7A and TTC7B there are multiple putative phosphorylated residues in this region (*36*), which may alter the conformation of the EFR3 binding surface. Further analysis of differences in EFR3 binding to TTC7A or TTC7B containing PI4KA complexes and how phosphorylation of TTC7A/B alters EFR3 binding will be required to understand the dynamic regulation of PI4KA membrane recruitment.

EFR3 proteins act as the membrane localization signal for the PI4KA complex (*5*), with a palmitoylation code controlling their spatial organisation between liquid-ordered and disordered regions of the PM (*23*). It has been previously revealed that there is an aspect of plasticity in how EFR3 recruits PI4KA, as the integral membrane protein TMEM150A can function in tandem with EFR3 to recruit PI4KA to the PM in the absence of TTC7-FAM126 (*23*, *37*). Our structure reveals the canonical mechanism of PM recruitment mediated by TTC7-FAM126. EFR3A mutations developed in this study that disrupt only TTC7-FAM126-mediated recruitment may also act as unique tools to discern between the diverse mechanisms of PI4KA recruitment to the PM. The triple helical binding interface of EFR3 that binds to TTC7-FAM126 contains multiple phosphorylation sites, present in both EFR3A and EFR3B (*36*). The location of these sites is divergent between EFR3A and EFR3B, potentially suggesting differential regulation (**Supplemental Fig 6B**). Intriguingly, the interfacial F728 in EFR3A is not conserved in EFR3B, with this residue being Y723 in EFR3B. This residue can be phosphorylated, with this PTM likely to dramatically decrease binding due to disruption of the TTC7-EFR3B interface. Further identification of the kinases and phosphatases that target these sites will be required to fully understand how the EFR3 interface can be regulated.

While subtle differences between complexes of TTC7A and TTC7B with the EFR3 C-terminus and FAM126 may exist, our structure of EFR3A bound to PI4KA-TTC7B-FAM126A still allows for interrogation of the possible molecular mechanisms underlying missense, nonsense and point mutations in TTC7A that cause severe gastrointestinal and immunological disease. This disease is an autosomal-recessive inherited disease, with patients having heterogeneous intestinal and immunologic disease presentations. The most severe TTC7A mutations cause nonsense or frameshift mutations in TTC7A that collectively remove both the EFR3 and PI4KA binding sites (*34*). However, there are numerous missense mutations that cluster to regions in spatial proximity to the putative EFR3 binding site in TTC7A. The mutations S539L, A551D, and H570R are all in helices that either bind the α3 helix of EFR3A, or pack against the interfacial helices, with these likely altering the orientation of the EFR3A binding site. In addition, K606R is predicted to pack against the helix 704-724 in TTC7A (equivalent to the EFR3 interfacial helix 690-707 in TTC7B). This mutation would likely alter the orientation of the EFR3 interface, causing decreased affinity. Further biochemical and biophysical analysis will be required to define the effect of disease-causing TTC7A mutations on EFR3-mediated membrane recruitment versus PI4KA complex stability.

A limitation of the molecular insight we have observed for how EFR3 interacts with the PI4KA complex is that all our biophysical observations on the association of the EFR3 C-terminus with the PI4KA complex were carried out exclusively in solution. The biologically relevant association of EFR3 will occur in the context of the full length EFR3 localised on a membrane surface. While we have no expectation of the ordered domains of EFR3 altering how the C-terminus binds to the PI4KA complex, there likely will be a critical role of this region in proper EFR3 membrane localisation. For the evolutionarily similar class I-III PI3Ks and PI4KB there are extensive conformational changes in the lipid kinase domain that are driven by association with membrane substrate, and it is likely that this will also be the case for PI4KA. To completely identify the full repertoire of regulatory mechanisms underlying PI4KA activation future experiments will need to address conformational dynamics of full-length lipidated EFR3 recruitment of the PI4KA complex in its native membrane environment.

Collectively, our detailed biochemical and structural analyses of the EFR3A-mediated recruitment of PI4KA complex to the PM provides unique insight into the assembly and regulation of PI4KA complexes. This work also provides a framework to further explore how PM recruitment of PI4KA is regulated. Overall, the findings from this study will be useful in developing novel therapeutic strategies to target aberrant PI4KA signaling in human disease.

## Acknowledgements

JEB is supported by the Canadian Institute of Health Research, and a Michael Smith Foundation for Health Research (Scholar Award 17686). ALS was supported by an NSERC CGS-M scholarship. C.K.Y. is supported by CIHR (PJT-168907) and the Natural Sciences and Engineering Research Council of Canada (RGPIN-2018-03951). Cryo-EM specimens were prepared, and data was collected at the High-Resolution Macromolecular Electron Microscopy (HRMEM) facility at the University of British Columbia (https://cryoem.med.ubc.ca). We thank Claire Atkinson, Joeseph Felt, Liam Worrall and Natalie Strynadka. HRMEM is funded by the Canadian Foundation for Innovation and the British Columbia Knowledge Development Fund. The work of TB, JGP, PR and AAP was funded by the Intramural Research Program of the *Eunice Kennedy Shriver* National Institute of Child Health and Human Development of the National Institutes of Health, Bethesda, MD, USA. (HHS/NIH/NICHD - Z01:HD000196-25). Confocal imaging was performed at the Microscopy & Imaging Core of the NICHD with the kind assistance of Drs. Vincent Schram and Ling Yi.

## Competing Interests

J.E.B. reports personal fees from Scorpion Therapeutics and Reactive therapeutics; and research contracts from Novartis and Calico Life Sciences.

## Data Availability

The EM data have been deposited in the EM data bank with accession number (EMDB: 44413), and the associated structural model has been deposited to the PDB with accession number (PDB: 9BAX). The MS proteomics data have been deposited to the ProteomeXchange Consortium via the PRIDE partner repository with the dataset identifier PXD043442 (*38*). All raw data in all figures are available in the source data excel file.

## Methods

### Plasmid constructs

Plasmids containing genes for full length PI4KA, TTC7B and FAM126A (1-308) were cloned into the MultiBac vector, pACEBac1 (Geneva Biotech) as described in (*21*). Plasmid encoding TTC7B and FAM126A (1-308) was subcloned into a pOPT vector containing an N-terminal 10X histidine tag, followed by a 2X Strep tag, followed by a Tobacco Etch Virus (TEV) protease cleavage site. Based on the evolutionary conservation of the C-terminal tail of EFR3A (**Fig. S1**), we designed and synthesized a construct containing the C-terminus of EFR3A from residue 688 to 821 from GeneArt gene synthesis (Invitrogen). EFR3A (721-791) was subcloned into two pOPT vectors; one containing an N-terminal 10X histidine tag and a 2X Strep tag, and one containing an N terminal Maltose Binding Protein (MBP) tag in addition to the 10X Histidine tag and 2X Strep tag (*39*). EFR3A, TTC7B and FAM126A substitution mutants shown in Table S2 were generated using site-directed mutagenesis according to published protocols using EFR3A (721-791) as template. Plasmid encoding human EFR3B^HA^-T2A-TTCB^MYC^-P2A-^FLAG^FAM126A in pcDNA3.1 vector, which is referred to more simply here as EFR3B-P2A-TTC7B-T2A-FAM126A was a kind gift from the Cyert lab at Stanford university. EFR3B, TTC7B and FAM126A substitution and deletion mutants shown in Table S2 in the pcDNA3.1 vector were generated using Gibson assembly (*39*).

Alternatively, the high-sensitivity PM-PI4KA^BRET^ biosensor (L10-mVenus-tPT2A-nLuc-PI4KA) was constructed in multiple steps by standard restriction cloning using enzymes from New England Biolabs. First, the sLuc cassette in the L10-mVenus-T2A-sLuc-(2X)P4M biosensor (*40*) was exchanged for NanoLuciferase (nLuc) using an insert amplified from PM-nLuc (Forward Primer, 5’-AT ATA CCG GTC ATG GTC TTC ACA CTC GAA GAT TTC GTT GG −3’; Reverse Primer, 5’-AT ATC TCG AGA CGC CAG AAT GCG TTC GCA CAG C −3’), which was a kind gift from Laszlo Hunyady (Addgene Plasmid # 164784), together with AgeI and XhoI restriction sites. Next, we replaced the single T2A site in the resulting L10-mVenus-T2A-nLuc-(2X)P4M biosensor with the tandem tPT2A sequence (*41*) using a custom-synthesized insert from GeneArt (Invitrogen) and a SalI and AgeI double-digest. Finally, we replaced the (2X)P4M domain in the L10-mVenus-tPT2A-nLuc-(2X)P4M vector with an in-frame PI4KA insert derived from pEGFP-PI4KA (*42*) using shared XhoI and MfeI restriction sites in the parent C1 vector backbones (CloneTech). Whole plasmid sequencing for all plasmids was performed by Plasmidsaurus using Oxford Nanopore Technology with custom analysis and annotation.

### Protein Expression

Plasmids containing the coding sequences for EFR3A (721-791), EFR3A mutants and MBP-EFR3A (721-791) were expressed in BL21 DE3 C41 *Escherichia coli* and induced with 0.5 mM IPTG and grown at 37 °C for 4 h. The TTC7B-FAM126A (1-308) construct was expressed in BL21 DE3 C41 cells and induced with 0.1mM IPTG and grown at 21 °C for 20 hours. Cells were then harvested and centrifuged at 15,000*g*. Pellets were washed with PBS before being snap-frozen in liquid nitrogen, followed by storage at −80 °C. Bacmid harboring PI4KA complex (PI4KA-TTC7B-FAM126A (1-308)) was transfected into *Spodoptera frugiperda (Sf9) cells*, and viral stocks amplified for one generation to acquire a P2 generation final viral stock. Final viral stocks were added to *Sf9* cells in a 1/100 virus volume to cell volume ratio. Constructs were expressed for 65-72 hours before harvesting of the infected cells. Cell pellets were washed with PBS, flash frozen in liquid nitrogen, and stored at −80 °C.

### Protein Purification

#### EFR3A, EFR3A mutants and MBP-EFR3A purification

Cell pellets were lysed by sonication for 5 min in lysis buffer (20 mM Tris [pH 8.0], 100 mM NaCl, 5% [v/v] glycerol, 20 mM imidazole, 2 mM β-mercaptoethanol [bME]), and protease inhibitors [Millipore Protease Inhibitor Cocktail Set III, EDTA free]). Triton X-100 was added to 0.1% v/v, and the solution was centrifuged for 45 min at 20,000*g* at 1 °C (Beckman Coulter J2-21, JA-20 rotor). The supernatant was then loaded onto a 5 ml HisTrap column (Cytiva) that had been equilibrated in nickel–nitrilotriacetic acid (Ni–NTA) A buffer (20 mM Tris [pH 8.0], 100 mM NaCl, 20 mM imidazole [pH 8.0], 5% [v/v] glycerol, and 2 mM bME). The column was washed with of high salt buffer (20 mM Tris [pH 8.0], 1 M NaCl, 5% [v/v] glycerol, and 2 mM bME), followed by of 6% Ni–NTA B buffer (20 mM Tris [pH 8.0], 100 mM NaCl, 200 mM imidazole [pH 8.0], 5% [v/v] glycerol, and 2 mM bME) before being eluted with 100% Ni–NTA B. The eluate was then loaded onto a 5 ml Strep column and then washed with GFB (20 mM HEPES [pH 7.5], 150 mM NaCl, 5% [v/v] glycerol and 1 mM Tris(2-carboxyethyl) phosphine [TCEP]). Protein was eluted with GFB containing 2.5 mM desthiobiotin and concentrated in a 10 kDa MWCO concentrator (Millipore Sigma). Concentrated protein was loaded onto the Superdex 75 Increase 10/300 GL (Cytiva) or the Superdex 200 Increase 10/300 GL (Cytiva) pre-equilibrated in GFB. Protein fractions were collected and concentrated in a 10 kDa MWCO concentrator (Millipore Sigma), flash frozen in liquid nitrogen, and stored at −80 °C.

#### TTC7B-FAM126A (1–308) purification

Cell pellets were lysed by sonication for 5 min in lysis buffer (20 mM imidazole [pH 8.0], 100 mM NaCl, 5% [v/v] glycerol, 2 mM bME), and protease inhibitors [Millipore Protease Inhibitor Cocktail Set III, animal-free]. Triton X-100 was added to 0.1% v/v, and the solution was centrifuged for 45 min at 20,000*g* at 1 °C. The supernatant was then loaded onto a 5 ml HisTrap column (Cytiva) that had been equilibrated in nickel–nitrilotriacetic acid (Ni–NTA) A buffer (20 mM Tris [pH 8.0], 100 mM NaCl, 20 mM imidazole [pH 8.0], 5% [v/v] glycerol, and 2 mM bME). The column was washed with Ni-NTA A buffer (20 mM imidazole [pH 8.0], 100 mM NaCl, 5% [v/v] glycerol, and 2 mM bME) followed by 6% Ni–NTA B buffer (450 mM Imidazole [pH 8.0], 100 mM NaCl, 5% [v/v] glycerol, and 2 mM bME) before being eluted with 100% Ni– NTA B. The eluate was then loaded onto a 5 ml Strep column and then washed with filtration buffer (GFB) (20 mM Imidazole [pH 7.0], 150 mM NaCl, 5% [v/v] glycerol and 0.5 mM Tris(2-carboxyethyl) phosphine (TCEP)). The column was loaded with tobacco etch virus protease containing a stabilizing lipoyl domain. TEV cleavage proceeded overnight following which the protein was eluted with GFB. Protein eluate was further concentrated in a 50 kDa MWCO concentrator (Millipore Sigma). Concentrated protein was loaded onto the Superdex 200 Increase 10/300 GL (Cytiva) or the Superose 6 Increase 10/300 GL (Cytiva) pre-equilibrated in GFB. Protein fractions from a single peak were collected and concentrated in 50 kDa MWCO concentrator (Millipore Sigma), flash frozen in liquid nitrogen and stored at −80 °C until further use. This protein is referred to as TTC7B-FAM126A.

#### PI4KA-TTC7B-FAM126A (1-308) purification

Sf9 pellets were resuspended in lysis buffer [20 mM imidazole pH 8.0, 100 mM NaCl, 5% glycerol, 2 mM βMe, protease (Protease Inhibitor Cocktail Set III, Sigma)] and lysed by sonication. Triton X-100 was added to 0.1% v/v final, and lysate was centrifuged for 45 min at 20,000×g at 1 °C. The supernatant was then loaded onto a 5 ml HisTrap column (Cytiva) that had been equilibrated in nickel–nitrilotriacetic acid (Ni–NTA) A buffer (20 mM Tris [pH 8.0], 100 mM NaCl, 20 mM imidazole [pH 8.0], 5% [v/v] glycerol, and 2 mM bME). The column was washed with Ni-NTA A buffer (20 mM imidazole [pH 8.0], 100 mM NaCl, 5% [v/v] glycerol, and 2 mM bME) followed by 6% Ni–NTA B buffer (27 mM Imidazole [pH 8.0], 100 mM NaCl, 5% [v/v] glycerol, and 2 mM bME) before being eluted with 100% Ni–NTA B. Elution fractions were passed through a 5 ml StrepTrapHP column (Cytiva) pre-equilibrated in Gel Filtration buffer (GFB) (20 mM imidazole pH 7.0, 150 mM NaCl, 5% glycerol [v/v], 0.5 mM TCEP). The column was washed with GFB before loading a tobacco etch virus protease containing a stabilizing lipoyl domain (Lip-TEV), and cleavage proceeded overnight. Cleaved protein was eluted with GFB and concentrated in a 50 kDa MWCO concentrator (MilliporeSigma). Concentrated protein was loaded onto the Superose 6 Increase 10/300 GL (Cytiva) pre-equilibrated in GFB. Protein fractions from a single peak were collected and concentrated in 50 kDa MWCO concentrator (Millipore Sigma), flash-frozen in liquid nitrogen and stored at −80 °C until further use. This protein is referred to as PI4KA complex.

The raw SDS-PAGE gel images for all purified proteins after gel filtration are shown in the source data.

### Co-elution of PI4KA complex-EFR3A and TTC7B-FAM126A (1-308)-EFR3A

7.19 μM of PI4KA complex and 61.8 μM of EFR3A were loaded onto a Superose 6 Increase column pre-equilibrated in GFB [150 mM NaCl, 20 mM imidazole (pH 7.0), 5% glycerol (v/v), 0.5 mM TCEP]. Fractions containing the PI4KA complex bound to EFR3A were pooled, concentrated, flash-frozen in liquid nitrogen and stored at −80°C. Protein was run on an SDS-PAGE gel to confirm the formation of the complex. For the co-elution of TTC7B-FAM126A (1-308) and EFR3A, we used 6.78 μM of TTC7B-FAM126A and 102.3 μM of EFR3A and performed size exclusion chromatography as mentioned above. The peaks were run on a SDS-PAGE gel to confirm formation of the complex.

### Protein purification for Cryo-EM

Pre-Gel filtered PI4KA complex (13.0 μM final), and gel filtered EFR3A (113 μM final) were incubated together on ice for 20 minutes prior to injection onto the Superose 6 Increase 10/300 GL (Cytiva) pre-equilibrated in GFB (150 mM NaCl, 20 mM imidazole [pH 7.0], 5% glycerol [v/v], 0.5 mM TCEP). Protein fractions from a single peak were collected and concentrated to 250 μl at 4μM and BS3 was added to a final concentration of 1 mM. The reaction was incubated on ice for 1 hour. 1 M Tris (pH 7.5) was added to a final concentration of 50 mM to quench the reaction and incubated at RT for 15 minutes. The quenched reaction was loaded onto the Superose 6 Increase 10/300 column pre-equilibrated in GFB. Protein fractions from the main peak, consistent with an elution volume of the PI4KA complex-EFR3A (721-791), were collected, concentrated, flash frozen in liquid nitrogen, and stored at −80°C.

### Cryo-EM sample preparation and data collection

C-Flat 2/1-3Cu-T-50 grids mesh grids were glow-discharged for 25 s at 15 mA using a Pelco easiGlow glow discharger. 3 μL of crosslinked PI4KA complex-EFR3A (721–791) were applied to the grids at 0.7 mg/ml. The grids were prepared using a Vitrobot Mark IV (FEI) by blotting for 1.5 seconds at a blot force of −5 at 4°C and 100% humidity before plunge freezing in liquid ethane. Grids were screened for particle and ice quality at the UBC High Resolution Macromolecular Cryo-Electron Microscopy (HRMEM) facility using a 200-kV Glacios transmission microscope (Thermo Fisher Scientific) equipped with a Falcon 3EC direct electron detector (DED). Datasets were collected at the UBC HRMEM facility using a 300-kV Titan Krios equipped with Selectris energy filter with a Falcon IV camera. A total of 9,412 super-resolution movies were collected using SerialEM with a total dose of 50 e^-^/Å^2^ over 774 frames at a physical pixel size of 0.77 Å per pixel, using a defocus range of −0.5 to −2 um, at 165,000 x magnification.

### Cryo-EM image processing

All data processing was carried out with cryoSPARC v4.2.1. Patch motion correction using default settings was applied to all movies to align the frames and Fourier-crop the outputs by a factor of 2. The contrast transfer function (CTF) of the resulting micrographs was estimated using the patch CTF estimation job with default settings. Micrographs were manually curated to contain micrographs with CTF fit resolution less than or equal to 10. To generate an initial model, 277,573 particles were picked from 3,501 micrographs using blob picking with a minimum and maximum diameter of 250 and 280, respectively. Particles were inspected using the inspect picks job to remove particles that picked ice contamination and were then extracted with a box size of 500 pixels, for a total of 201,081 particles. The particles were subjected to 2D classification using default settings. The best class averages were then selected and used as an initial model for template picking. These 2D classes were low pass filtered to 20 Å and used to template pick with the particle diameter set to 300 Å. Particles were inspected and 510,887 were extracted with a box size of 600 pixels and subjected to two rounds of 2D classification with 40 online-EM iterations to remove particles with ice contamination or those that showed no features. A total of 77,624 particles were used for ab initio reconstruction and heterogeneous refinement using three classes, with C1 symmetry. Given the previously solved PI4KA/TTC7B/FAM126A (2-289) cryo-EM structure employing C2 symmetry (*18*), the following homogenous refinements were done with C2 symmetry. The best heterogeneous refinement which had 46,039 particles underwent two rounds of homogenous refinement. These 46,039 particles were subjected to further ab initio 3D reconstructions, heterogeneous refinement, and homogeneous refinement jobs to filter out incorrectly assigned particles. The final initial model to be used to generate templates for auto-picking of particles consisted of 8,647 particles.

To generate a cryo-EM map from the complete dataset, the collected movies were split into three groups (Group 1: 3,535 movies, Group 2: 3,364 movies, Group 3: 2,513 movies) and processed into micrographs as described above. The initial template from above was used to auto-pick 493,392 (group 1), 415,293 (group 2), and 307,632 (group 3) particles, using the template picker job with particle diameter set to 300 Å and a minimum separation distance of 0.75 diameters. Particles were inspected as above and 275,667 (group 1), 264,196 (group 2), and 185,766 (group 3) particles were extracted with a box size of 600 pixels. Each group of particles independently underwent one (group 1) or two (groups 2 and 3) rounds of 2D classification with 40 online-EM iterations. The resulting 105,657 (group 1), 57,805 (group 2), and 77,900 (group 3) particles were independently used for ab initio reconstruction using 3 classes and subsequent heterogeneous refinement with C1 symmetry. The best 3D classification from groups 1 and 2 were merged into 93,029 particles (55,802 (group 1) + 37,227 (group 2)) and underwent two rounds of ab initio reconstruction with 3 classes followed by heterogenous refinement. The best 3D classification, consisting of 44,305 particles was refined using homogeneous refinement using C1 symmetry. The particles from group 3 were then added to these 44,305 particles. A total of 67,563 particles were used to carry out homogeneous refinement, followed by masked local refinement with a static mask using the previous 3D reconstruction as the starting model, yielding a reconstruction with an overall resolution of 3.48 Å based on the Fourier shell correlation (FSC) 0.143 criterion. The full workflow to generate the final cryo-EM map from the three groups is shown in Supplemental Fig 2.

#### AlphaFold3 modelling

We used the protein prediction software, AlphaFold3 (*43*) to predict where EFR3A’s c-terminus interacts with the TTC7B/FAM126A. Specifically, we used the AlphaFold server (https://golgi.sandbox.google.com/) and input the sequence for full length human TTC7B and FAM126A with the C-terminus of human EFR3A (aa 621-821). To evaluate the confidence of individual subunit assembly predictions, we analysed the predicted alignment error (pae), predicted template modelling score (PTM) and the interface predicted template modelling score. The top ranked model had ptm and iptm scores of 0.73 and 0.79, respectively, consistent with a stable complex. The chain_pair_iptm scores are useful in evaluating the confidence of predicted protein-protein interfaces. The chain_pair_iptm score for TTC7B:FAM126A was 0.82, TTC7B:EFR3A was 0.63, and FAM126A:EFR3A was 0.35. The chain_pair_pae_min values correlate with whether two chains interact with each other. The chain_pair_pae_min score for TTC7B:FAM126A was 1.15, TTC7B:EFR3A was 1.88, and FAM126A:EFR3A was 1.6. These iptm and pae scores are consistent with a high confidence protein complex prediction.

### Model building

The cryo-EM structure of PI4KA-TTC7B-FAM126A (PDB: 6BQ1) (*18*) was fit into the map using Chimera(*44*). The 1^st^ helix of EFR3A was at sufficient local resolution for iterative rounds of automated model building in Phenix, manual model building in COOT, and refinement in Phenix.real_space_refine using realspace, rigid body, and adp refinement with tight secondary structure restraints (*45*). For the remainder of the EFR3A interface with FAM126A and TTC7B we used AlphaFold-Multimer using Google Colab (*28*)(https://colab.research.google.com/github/sokrypton/ColabFold/blob/main/AlphaFold 2.ipynb) was used to model the structure of TTC7B-FAM126A–EFR3A. The sequences of full length TTC7B, residues 1-360 of FAM126A with residues 681-821 of EFR3A were input to generate the model of the complex. AlphaFold generated five models ranked in order by mean pLDDT. The highest ranked model showed strong correlation to the cryo-EM density, and was used to model the 2^nd^ and 3^rd^ helix of EFR3A bound to FAM126A and TTC7B, with multiple rounds of refinement in Phenix.real_space_refine using realspace, rigid body, and adp refinement with tight secondary structure restraints (*45*). The full refinement and validation statistics are shown in **Supplementary Table 1**.

### HDX-MS sample preparation

HDX reactions examining the TTC7B-FAM126A (1-308) in the presence and absence of EFR3A were carried out in 15 µL reaction volumes containing 15 pmol of protein (1 µM TTC7B-FAM126A, 1 µM EFR3A in the bound state). The exchange reactions were initiated by the addition of 12.55 µL of D_2_O buffer (20 mM Imidazole pH 7, 150 mM NaCl) to 2.45 µL of protein (final D_2_O concentration of 78.2% [v/v]). Reactions proceeded for 3s, 30s, 300s, 3000s, and 10,000s at 20°C before being quenched with ice cold acidic quench buffer, resulting in a final concentration of 0.6M guanidine HCl and 0.9% formic acid.

HDX reactions examining the PI4KA complex in the presence and absence of EFR3A were carried out in 15 µL reaction volumes containing 10 pmol of protein (0.667 µM PI4KA complex, 0.667 µM EFR3A in the bound state). The exchange reactions were initiated by the addition of 10.92 µL of D_2_O buffer (20 mM Imidazole pH 7, 150 mM NaCl) to 4.08 µL of protein (final D_2_O concentration of 68.0 % [v/v]). Reactions proceeded for 30s, 300s, 3000s, and 10,000s at 20°C before being quenched with ice cold acidic quench buffer, resulting in a final concentration of 0.6M guanidine HCl and 0.9% formic acid post quench.

HDX reactions examining MBP-EFR3A in the presence and absence of the TTC7B-FAM126A (1-308) were carried out in 20 µL reaction volumes containing 25 pmol of protein (1.25 µM MBP-EFR3A, 1.25 µM TTC7B-FAM126A in the bound state). The exchange reactions were initiated by the addition of 14.66 µL of D_2_O buffer (20 mM Imidazole pH 7, 150 mM NaCl) to 5.34 µL of protein (final D_2_O concentration of 68.5 % [v/v]). Reactions proceeded for 3s, 30s, and 300s at 20°C and 0.3s (3s on ice) before being quenched with ice cold acidic quench buffer, resulting in a final concentration of 0.6M guanidine HCl and 0.9% formic acid post quench.

All conditions and timepoints were created and run in independent triplicate. All samples were flash frozen immediately after quenching and stored at −80°C.

### Protein digestion and MS/MS data collection

Protein samples were rapidly thawed and injected onto an integrated fluidics system containing a HDx-3 PAL liquid handling robot and climate-controlled (2°C) chromatography system (Trajan), a Dionex Ultimate 3000 UHPLC system, as well as an Impact HD QTOF Mass spectrometer (Bruker). The full details of the automated LC system are described in (*46*). The TTC7B-FAM126A and PI4KA complex samples were run over one immobilized pepsin column (Trajan; ProDx protease column, 2.1 mm x 30 mm PDX.PP01-F32) at 200 µL/min for 3 minutes at 10°C, and the MBP-EFR3A samples were run over two immobilized pepsin columns (Waters; Enzymate Protein Pepsin Column, 300Å, 5µm, 2.1 mm X 30 mm) at 350 µL/min for 3 minutes at 2°C. The resulting peptides were collected and desalted on a C18 trap column (Acquity UPLC BEH C18 1.7mm column (2.1 mm x 5 mm); Waters 186004629). The trap was subsequently eluted in line with an ACQUITY 1.7 μm particle, 100 mm × 2.1 mm C18 UPLC column (Waters; 186003686), using a gradient of 3-35% B (Buffer A 0.1% formic acid; Buffer B 100% acetonitrile) over 11 minutes immediately followed by a gradient of 35-80% over 5 minutes. Mass spectrometry experiments acquired over a mass range from 150 to 2200 m/z using an electrospray ionization source operated at a temperature of 200°C and a spray voltage of 4.5 kV.

### Peptide identification

Peptides were identified from non-deuterated samples using data-dependent acquisition following tandem MS/MS experiments (0.5s precursor scan from 150-2000 m/z; twelve 0.25s fragment scans from 150-2000 m/z). TTC7B-FAM126A (1–308) and PI4KA complex MS/MS datasets were analysed using PEAKS7 (PEAKS), and peptide identification was carried out using a false discovery-based approach, with a threshold set to 1% using a database of purified proteins and known contaminants (*47*). The search parameters were set with a precursor tolerance of 20 ppm, fragment mass error 0.02 Da charge states from 1-8, leading to a selection criterion of peptides that had a −10logP score of 25.2 and 20.4 respectively. MBP-EFR3A (721–791) MS/MS datasets were analysed using FragPipe v18.0 and peptide identification was carried out by using a false discovery-based approach using a database of purified proteins and known contaminants (*47*, *48*). MSFragger was used, and the precursor mass tolerance error was set to −20 to 20ppm. The fragment mass tolerance was set at 20ppm. Protein digestion was set as nonspecific, searching between lengths of 4 and 50 aa, with a mass range of 400 to 5000 Da.

### Mass Analysis of Peptide Centroids and Measurement of Deuterium Incorporation

HD-Examiner Software (Sierra Analytics) was used to automatically calculate the level of deuterium incorporation into each peptide. All peptides were manually inspected for correct charge state, correct retention time, appropriate selection of isotopic distribution, etc. Deuteration levels were calculated using the centroid of the experimental isotope clusters. Results are presented as relative levels of deuterium incorporation, and the only control for back exchange was the level of deuterium present in the buffer (78.2%, 68.0%, or 68.5%). Differences in exchange in a peptide were considered significant if they met all three of the following criteria: ≥5% change in exchange, ≥0.45 Da difference in exchange, and a p value <0.01 using a two tailed student t-test. The entire HDX-MS dataset with all the values and statistics are provided in the source data. Samples were only compared within a single experiment and were never compared to experiments completed at a different time with a different final D_2_O level. The data analysis statistics for all HDX-MS experiments are provided in the source data according to the guidelines of (*31*). HDX-MS proteomics data generated in this study have been deposited to the ProteomeXchange Consortium via the PRIDE partner repository with the dataset identifier PXD043442 (*38*).

### Bio-layer interferometry (BLI)

The BLI measurements were conducted using a Fortebio (Sartorius) K2 Octet using fiber optic biosensors. Anti–penta-His biosensors were loaded using purified MBP-EFR3A, which had a 10x His tag on the N-terminus. The biosensor tips were preincubated in the BLI buffer (20 mM HEPES [pH 7.5], 150 mM NaCl, 0.01%, bovine serum albumin, and 0.002% Tween-20) for 10 min before experiments began. The sequence of steps in each assay was regeneration, custom, loading, baseline, association, and dissociation. Every experiment was done at 25 °C with shaking at 1000 rpm. Technical replicates were performed by using the same fiber tip and repeating the steps outlined previously. Regeneration was performed by exposing the tips to regeneration buffer (Glycine pH 1.5) for 5s and BLI buffer for 5s and repeating the exposure for 6 cycles. BLI buffer was used for the custom, baseline, and dissociation steps; these steps were performed in the same well for a given sample. For the dose response in **Figure 3A**, MBP-EFR3A was diluted in BLI buffer to 200 nM and was loaded onto the anti penta-His biosensor tips. TTC7B-FAM126A was also diluted in BLI buffer from 900 nM to 9.2 nM and added to the appropriate association wells. Non-specific association was controlled by loading 200 nM of His-MBP onto the biosensor tips and subtracting its responses from the responses measured with MBP-EFR3A. The *K*_D_ was estimated by one site specific non-linear regression analysis on GraphPad Prism 7.0 for Mac OS X, Graphpad Software, www.graphpad.com.

For the mutant binding experiments, EFR3A was diluted in BLI buffer to 200 nM and TTC7B-FAM126A was diluted in BLI buffer to 500 nM. All BLI experiments were performed using two or three technical replicates. Means ± SD were used to present values. Statistical analysis between conditions was performed using an unpaired t-test assuming unequal variances analysing mutant responses to wildtype responses measured simultaneously during each assay. The concentration of a given mutant was compared with the same concentration for wild-type binding. The following legends are used for statistical significance: ***p<0.0001, **p<0.001, *p< 0.01 and ns p>0.01.

### Multiple Sequence Alignments

Sequences were aligned using Clustal Omega Multiple Sequence Alignment, and the aligned sequences were subsequently analysed by ESPript 3.0 (https://espript.ibcp.fr) to visualize conserved regions (*49*). The UniProt accession codes for the aligned sequences used for TTC7 in **Figure 3C** are Q86TV6 *(Homo sapiens)*, Q9ULT0 *(Homo sapiens)*, E9Q6P5 *(Mus musculus)*, A0A8J1LRC8 *(Xenopus laevis)*, A0A8M2B4X5 *(Danio rerio),* A0A0B4K7H0 *(Drosophila melanogaster)*. The UniProt accession codes for the aligned sequences used for FAM126 in **Figure 4B** are Q9BYI3 *(Homo Sapiens)*, Q8IXS8 *(Homo Sapiens)*, Q6P9N1 *(Mus musculus)*, Q6P121 *(Danio rerio)*, Q7K1C5 *(Drosophila melanogaster)*, Q6A586 *(Caenorhabditis elegans)*. The UniProt accession codes for the aligned sequences used for EFR3 in **Figures 3B, 4A and Supplemental Figure 1** are Q14156 *(Homo Sapiens)*, Q9Y2G0 *(Homo sapiens)*, Q8BG67 *(Mus musculus)*, Q641A2 *(Xenopus laevis)*, Q5SPP5 *(Danio rerio)*, Q8IGJ0 *(Drosophila melanogaster)*, Q09263 *(Caenorhabditis elegans)*, O59817 *(Schizosaccharomyces pombe)*, Q03653 *(Saccharomyces cerevisiae)*.

### Mammalian Cell Culture

HEK293A (Invitrogen) cells were cultured in Dulbecco’s Modified Eagle Medium (DMEM-high glucose; Gibco) containing 10% (vol/vol) FBS and supplemented with a 1% solution of penicillin/streptomycin (Gibco). Cells were maintained at 37°C and 5% CO_2_ in a humidified atmosphere and regularly tested for *Mycoplasma* contamination using a commercially available detection kit (InvivoGen). After thawing, cell cultures are also treated with plasmocin (InvivoGen) at 500 µg/mL for the initial three passages (6-9 days) as well as supplemented with 5 µg/mL of the prophylactic for all subsequent passages.

### Live-Cell Measurements using the PM-PI4KA^BRET^ Biosensor

HEK293A cells (0.75×10^5^ cells/well) were seeded in a 200 μL total volume to white-bottom 96 well plates pre-coated with 0.01% poly-L-lysine solution (Sigma) and cultured overnight. Cells were then transfected with 0.5 μg of the PM-PI4KA^BRET^ biosensor (L10-mVenus-tPT2A-nLuc-PI4KA) using Lipofectamine 2000 (1 μL/well). Lipofection was done within Opti-MEM (40 μL/well) according to the manufacturer’s protocol, but with the slight modification of removing the media containing the Lipofectamine-complexed DNA and replacing it with complete culture medium at between 4-6 hrs post-transfection. Where indicated, increasing amounts (100, 500, or 1000 ng/well) of specified plasmids consisting of either an empty vector (pcDNA3.1), wild-type positive control (EFR3B-P2A-TTC7B-T2A-FAM126A), or the indicated mutants of the PI4KA complex components (EFR3B^△cterm^-P2A-TTC7B-T2A-FAM126A, EFR3B^L726A,F751A^-P2A-TTC7B-T2A-FAM126A, EFR3B-P2A-TTC7B^A702R^-T2A-FAM126A, or EFR3B-P2A-TTC7B-T2A-FAM126A^A103R^) were co-transfected (0.5-1.5 μg/well total). BRET measurements were made at 37°C using a Tristar2 LB 942 Multimode Microplate Reader (Berthold Technologies) with customized emission filters (540/40 nm and 475/20 nm). Between 20-24 hrs post-transfection, the cells were quickly washed before being incubated for 30 mins in 50 µL of modified Krebs-Ringer buffer (containing 120 mM NaCl, 4.7 mM KCl, 2 mM CaCl_2_, 0.7 mM MgSO_4_, 10 mM glucose, 10 mM HEPES, and adjusted to pH 7.4) at 37°C in a CO_2_-independent incubator. After the pre-incubation period, the cell-permeable luciferase substrate, coelenterazine h (40 µL, final concentration 5 µM), was added and the signal from the mVenus fluorescence and nLuc luminescence were recorded using 485 and 530 nm emission filters over a 45 min baseline BRET measurement (90 sec / cycle). Detection time was always 500 ms for each wavelength. To ensure a stable baseline is achieved following the substrate addition, BRET measurements are presented as the average basal BRET signal over the final 30 minutes measurement interval (15-45 minutes). All measurements were carried out in triplicate wells and repeated in three independent experiments. From each well, the BRET ratio was calculated by dividing the 530 nm and 485 nm intensities, which were then normalized to an internal measurement of wells transfected with the PM-PI4KA^BRET^ biosensor alone.

### Live-cell Confocal Microscopy of EGFP-PI4KA Localization

HEK293A cells (2.5×10^5^ cells/dish) were plated with a final volume of 1.5 mL on 29 mm circular glass-bottom culture dishes (#1.5; Cellvis) pre-coated with 0.01% poly-L-lysine solution (Sigma). Cells were allowed to attach overnight prior to transfection with plasmid DNAs (0.5-0.75 μg/well) using Lipofectamine 2000 (2-5 μL/well; Invitrogen). Lipofection was done using small volumes of Opti-MEM (200 μL; Invitrogen) according to the manufacturer’s instructions, once again with the slight modification of removing the media containing the Lipofectamine-complexed DNA at 4-6 hrs post-transfection and replacing it with complete DMEM. After 18-20 hrs of transfection, cells were incubated in 1 mL of modified Krebs-Ringer solution (containing 120 mM NaCl, 4.7 mM KCl, 2 mM CaCl_2_, 0.7 mM MgSO_4_, 10 mM glucose, 10 mM HEPES, and adjusted to pH 7.4) and images were acquired at room temperature using a Zeiss LSM 880 (63x/1.40 N.A. Plan-Apochromat Oil DIC M27 Objective) laser-scanning confocal microscopes (Carl Zeiss Microscopy). Image acquisition was performed using the ZEN software system (Carl Zeiss Microscopy), while image preparation and analysis was done using the open-source FIJI platform(*50*).

## Supplementary Figures and Tables

**Supplemental Figure 1:**
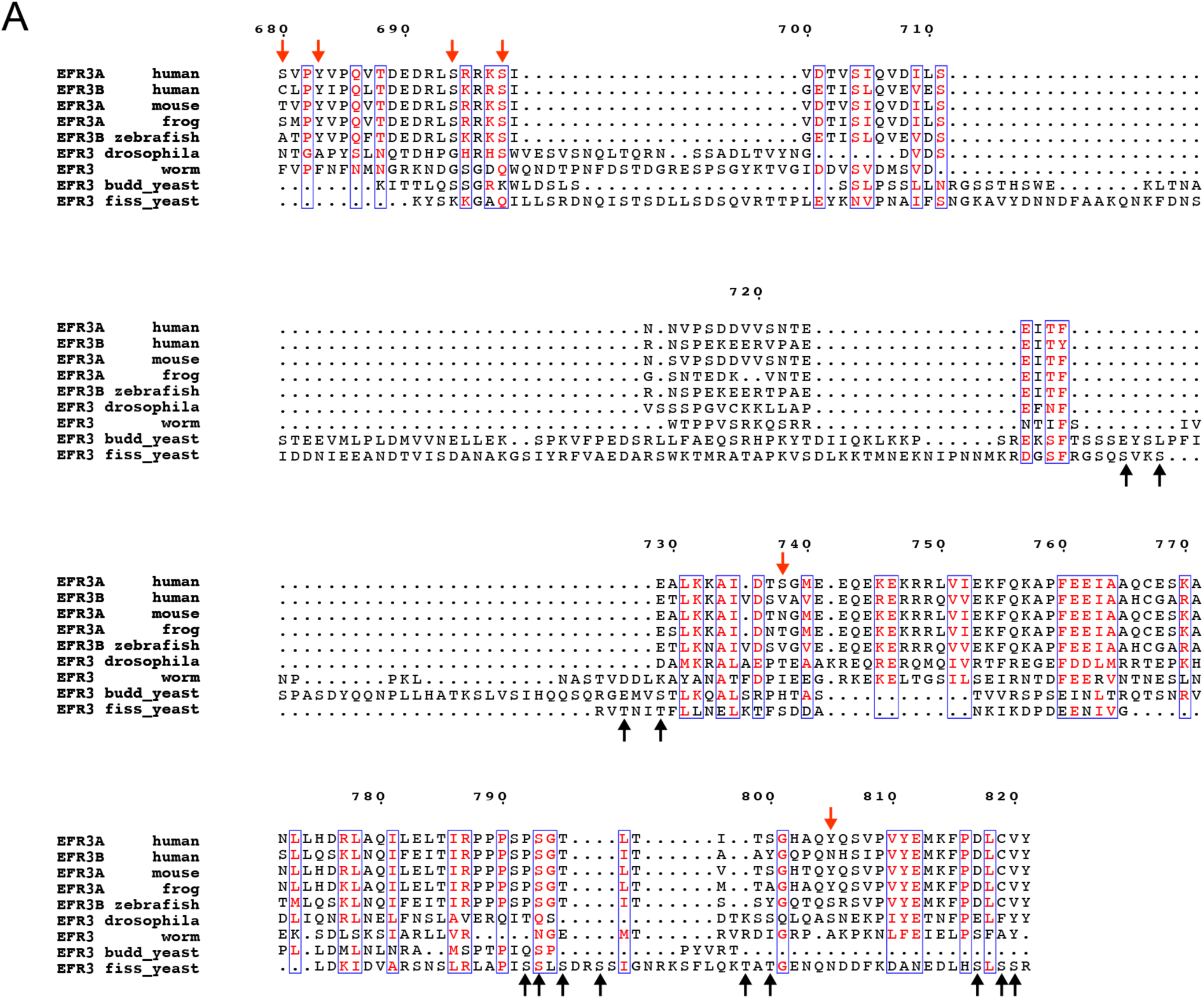
EFR3 C-terminal tail sequence alignment. **A.** Multiple sequence alignment (generated with ESPript 3.0) of EFR3 from *Homo sapiens, Mus musculus, Xenopus laevis, Danio rerio, Drosophila melanogaster, Caenorhabditis elegans*, *Schizosaccharomyces pombe,* and *Saccharomyces cerevisiae.* Human EFR3A residues that are reported phosphorylation sites (Phosphosite) are annotated above in red. Yeast efr3 residues that are reported phosphorylation sites (PhosphoGRID) are annotated below in black.

**Supplemental Figure 2 (related to Fig.1):**
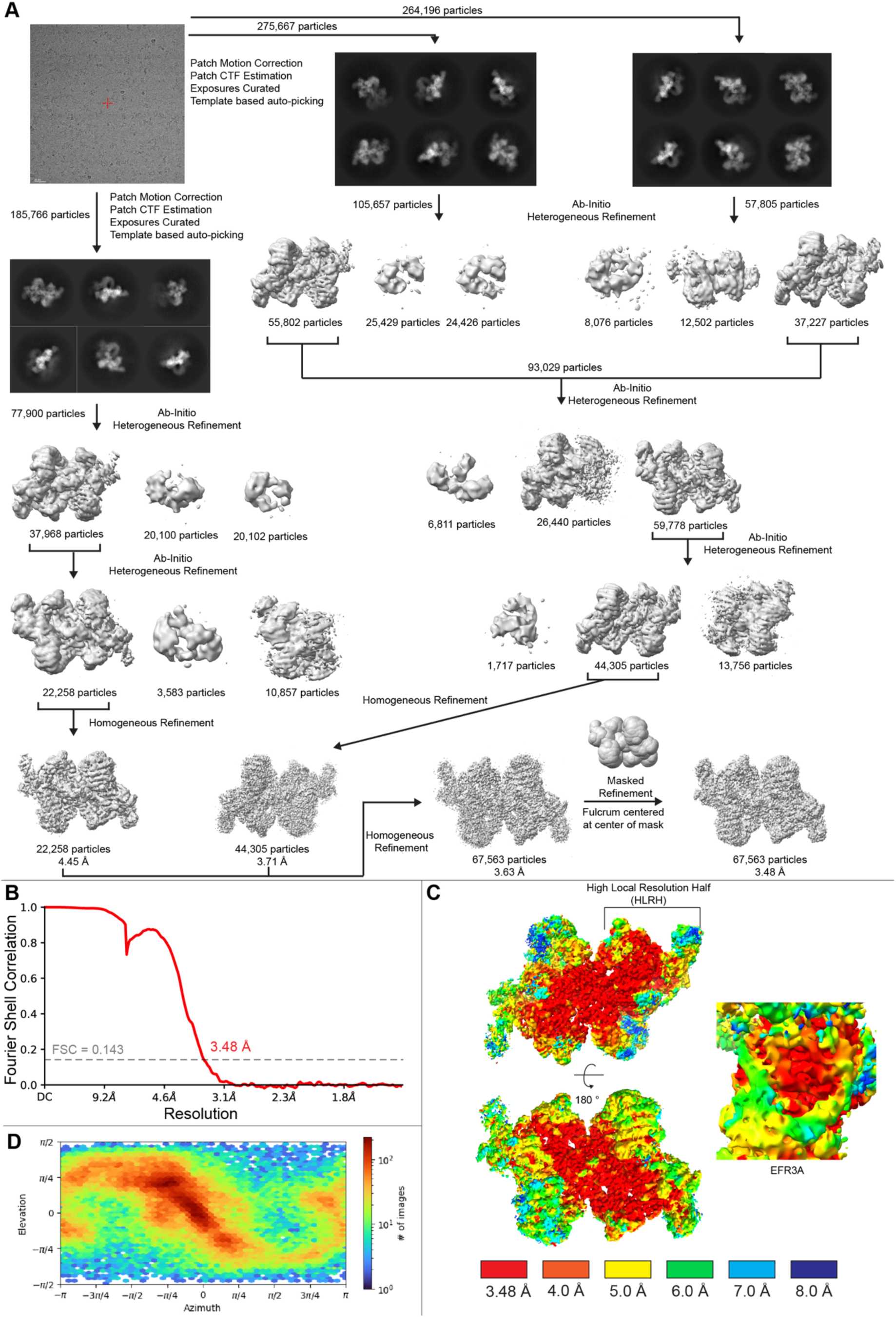
Cryo-EM data processing. **A.** Cryo-EM data processing workflow showing a representative micrograph from screening on the 200 kV Glacios, representative 2D class averages, and the image processing strategy used to generate a 3D reconstruction of the PI4KA/TTC7B/FAM126A/EFR3A complex. **B.** Gold standard Fourier shell correlation coefficient (FSC) curve after auto tightening by cryoSPARC for the final map. **C.** Final map coloured according to local resolution estimated using cryoSPARC v4.2.1, with a zoom-in of the EFR3A density. **D.** Viewing direction distribution plot of particles in the final cryo-EM reconstruction output by cryoSPARC v4.2.1.

**Supplemental Figure 3:**
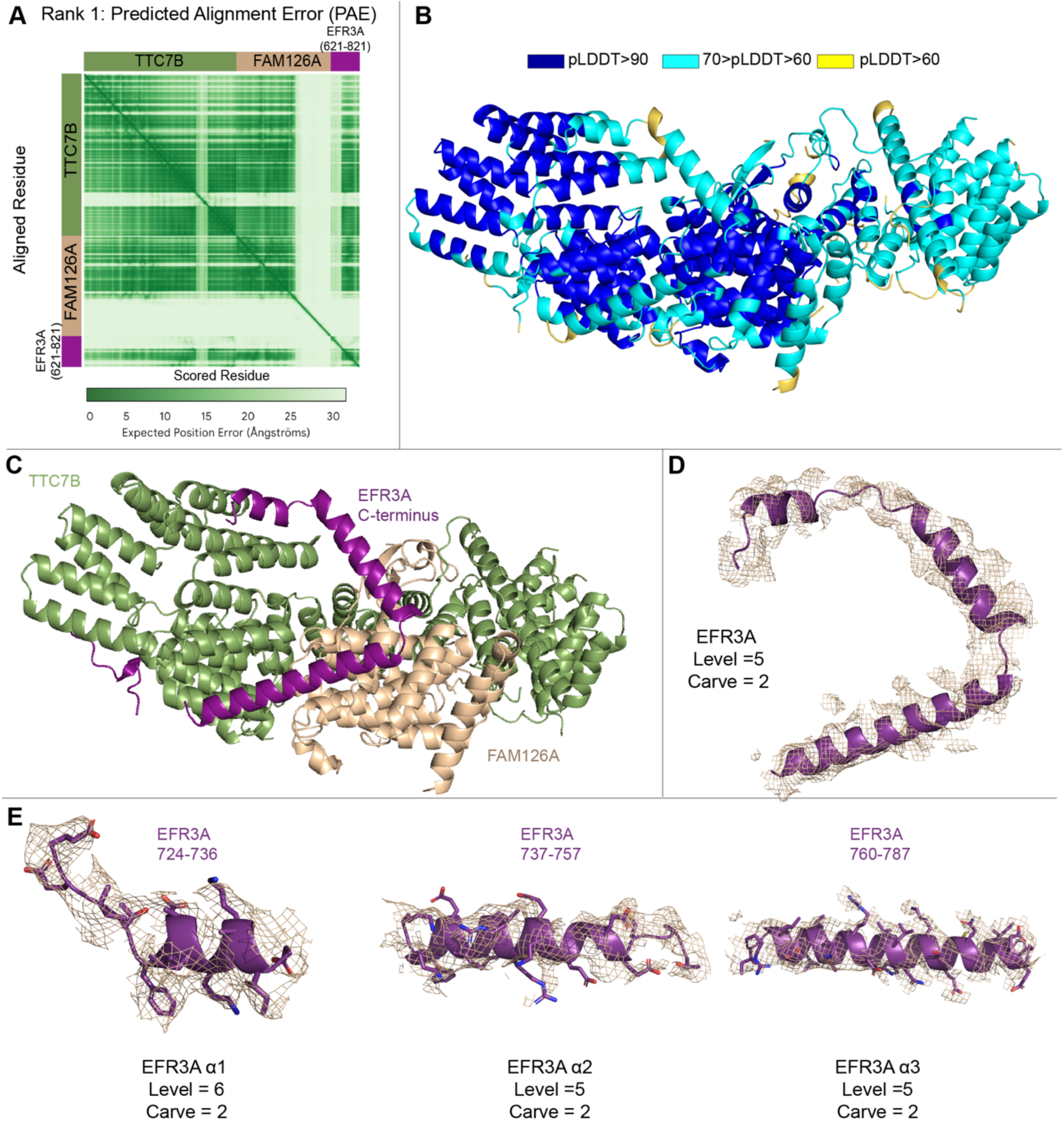
Model building with AlphaFold Multimer and map to model density fit. **A.** Predicted aligned error (PAE) of the AlphaFold3 prediction of the TTC7B-FAM126A and EFR3A C-terminus. **B.** AlphaFold3 model of TTC7B-FAM126A in complex with the EFR3A C-terminus with the per-residue confidence metric predicted local-distance difference test (pLDDT) <60 removed, coloured according to pLDDT score. **D.** AlphaFold3 model of the TTC7B-FAM126A in complex with EFR3A C-terminal with the per-residue confidence metric predicted local-distance difference test (pLDDT) <60 removed. **E.** Electron density of EFR3A C-terminus. **F.** Electron density of selected regions. (L) EFR3A α1 with density for EFR3A (724-736) shown on top, and (M) EFR3A α2 with density for EFR3A (737-759), and (R) EFR3A α3 with density for EFR3A (760-787) shown.

**Supplemental Figure 4:**
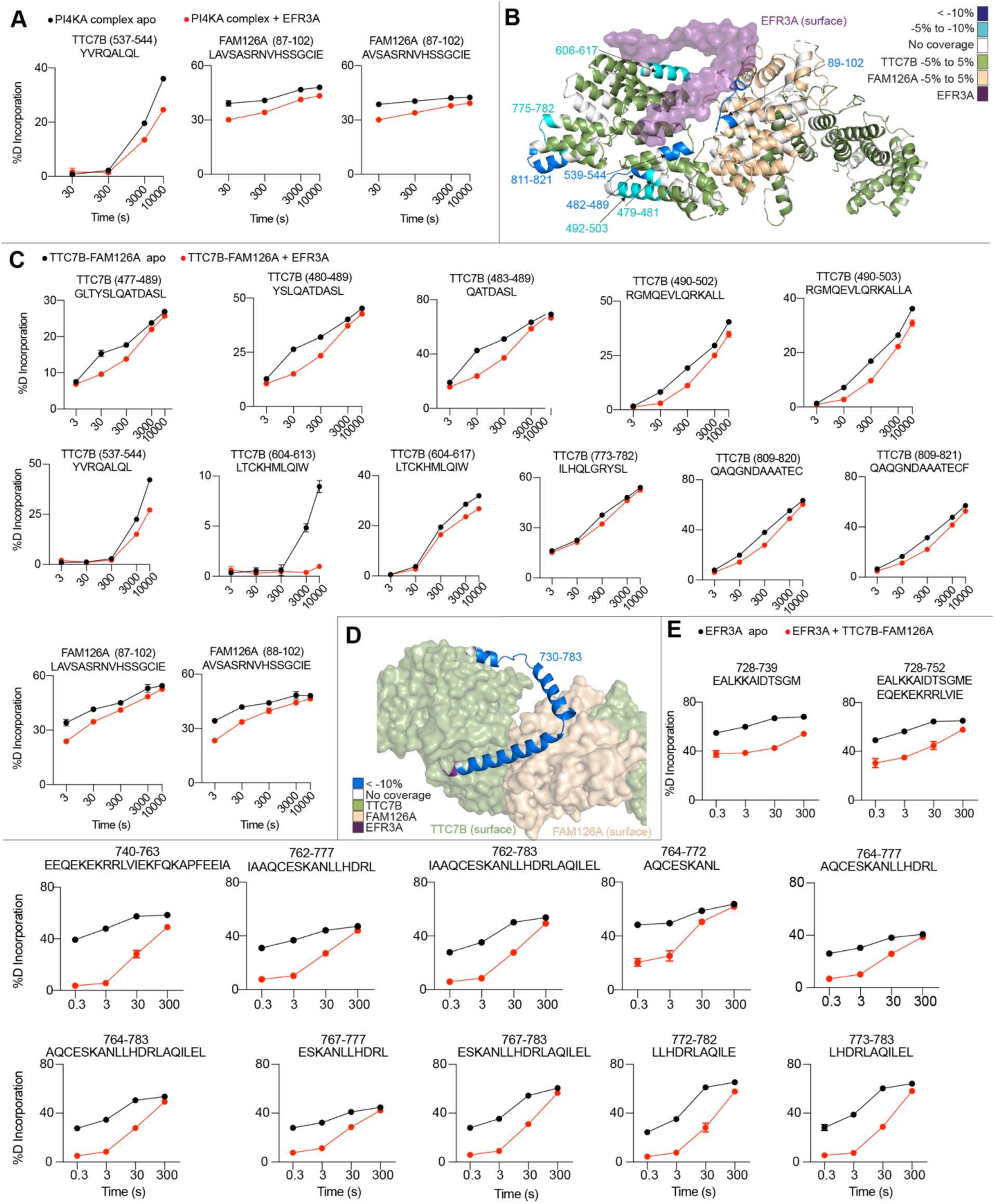
Raw HDX-MS deuterium incorporation curves. **A.** Relative deuterium incorporation traces for peptides showing significant differences in deuterium exchange (defined as >5% >0.45 Da, and p<0.01 in an unpaired two-tailed t-test at any time point) upon PI4KA complex binding to EFR3A. **B.** Significant differences in deuterium exchange (defined as >5% >0.45 Da, and p<0.01 in an unpaired two-tailed t-test at any time point) upon TTC7B-FAM126A dimer binding to EFR3A mapped on the high resolution half of our structural model with all regions annotated. **C.** Relative deuterium incorporation traces for peptides showing significant differences in deuterium exchange (defined as >5% >0.45 Da, and p<0.01 in an unpaired two-tailed t-test at any time point) upon TTC7B-FAM126A dimer binding to EFR3A. **D.** Significant differences in deuterium exchange (defined as >5% >0.45 Da, and p<0.01 in an unpaired two-tailed t-test at any time point) upon MBP-EFR3A binding to the TTC7B-FAM126A dimer mapped on the high resolution half of our structural model with all regions annotated. **E.** Relative deuterium incorporation traces for peptides showing significant differences in deuterium exchange (defined as >5% >0.45 Da, and p<0.01 in an unpaired two-tailed t-test at any time point) upon MBP-EFR3A binding to the TTC7B-FAM126A dimer.

**Supplemental Figure 5:**
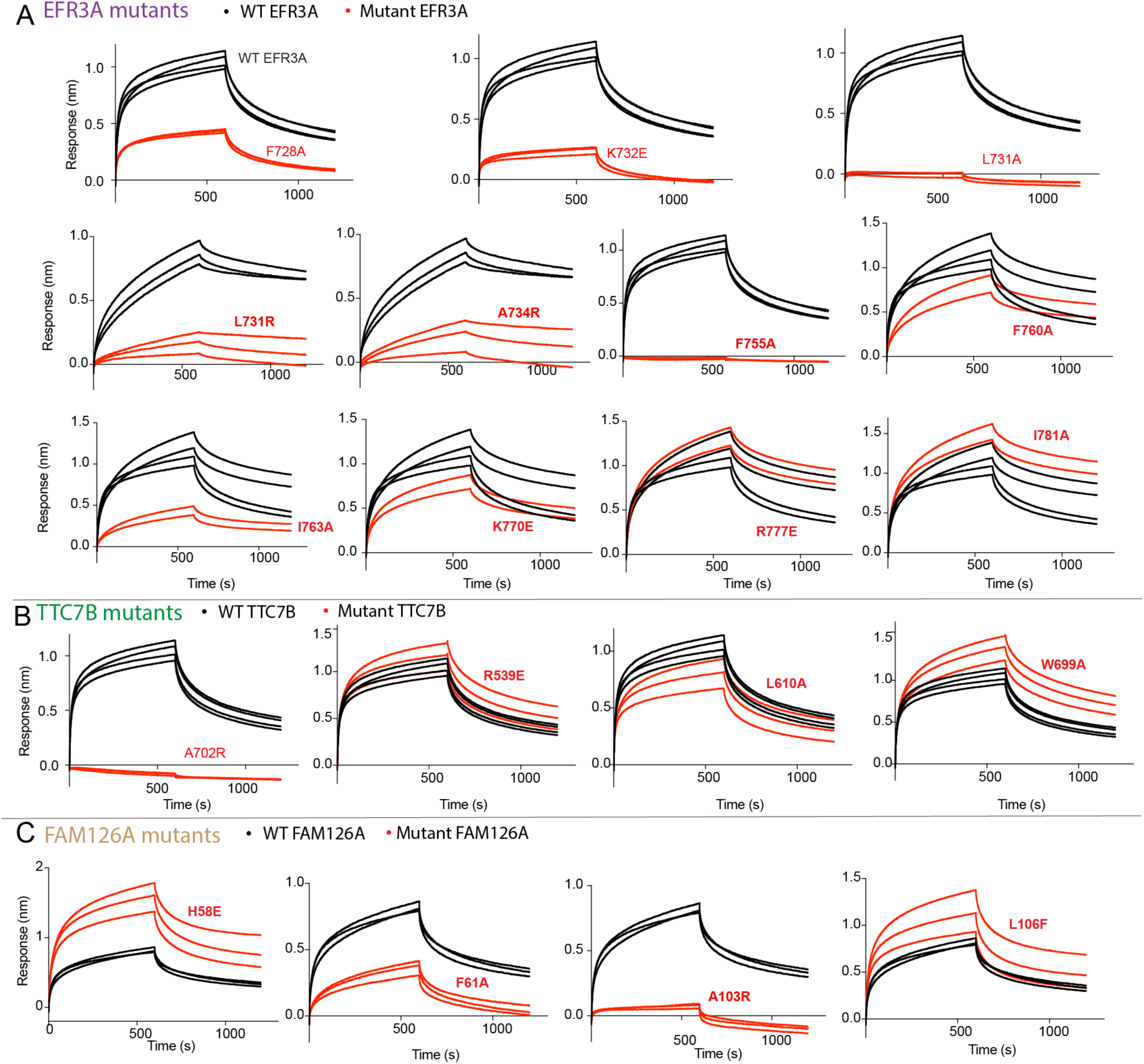
BLI raw data. **A.** Raw BLI association and dissociation curves of all EFR3A mutants (shown in table S2) compared to WT. His-EFR3A was loaded on the anti-penta-His tip at 200 nM and dipped in TTC7B-FAM126A at 500 nM. **B.** Raw BLI association and dissociation curves of all TTC7B mutants (shown in table S2) compared to WT. His-EFR3A was loaded on the anti-penta-His tip at 200 nM and dipped in TTC7B-FAM126A at 500 nM. **C.** Raw BLI association and dissociation curves of all TTC7B mutants (shown in table S2) compared to WT. His-EFR3A was loaded on the anti-penta-His tip at 200 nM and dipped in TTC7B-FAM126A at 500 nM.

**Supplemental Figure 6:**
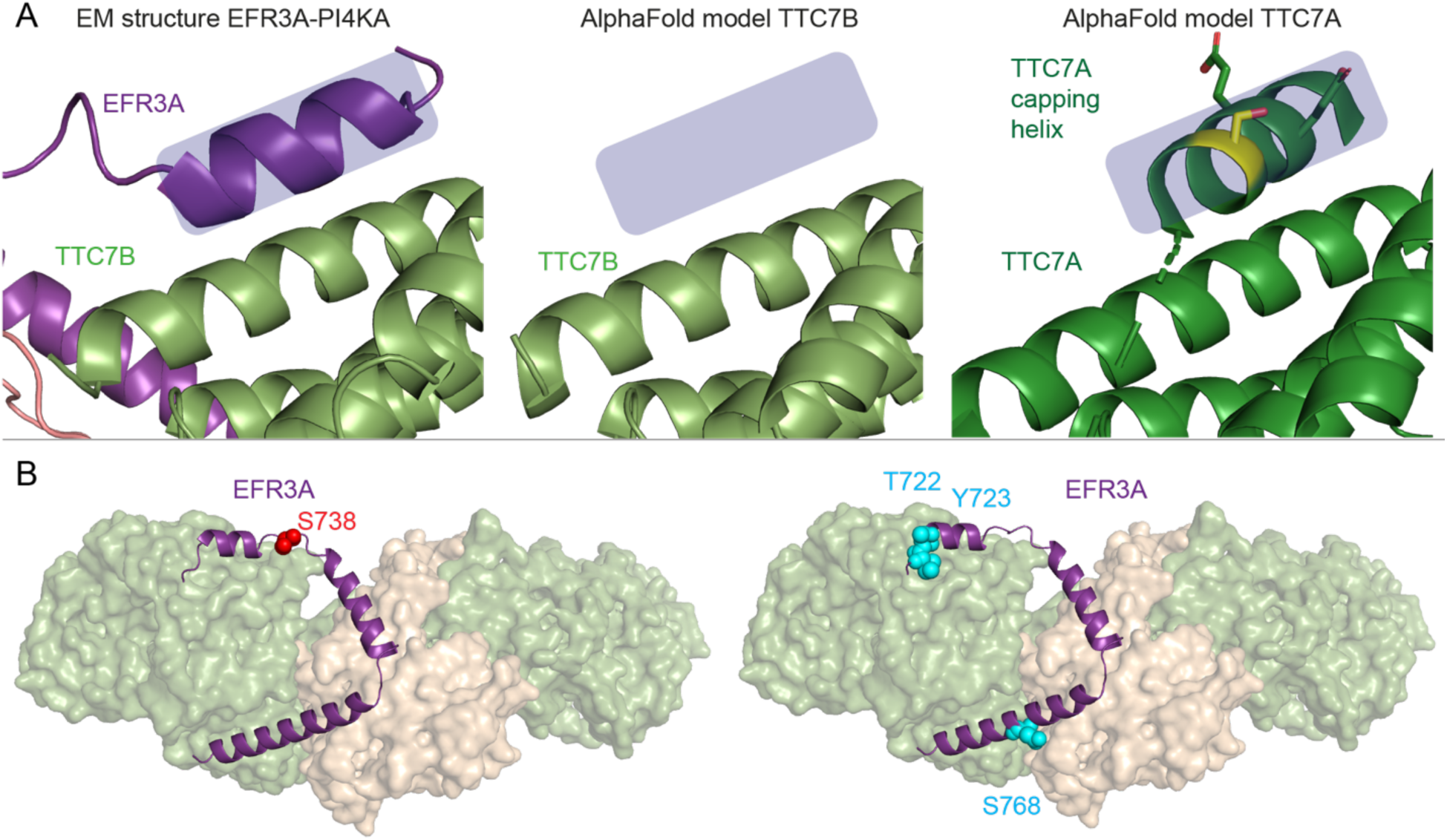
Differences between TTC7A/TTC7B and post-translational modification sites present at the EFR3-TTC7-FAM126 interface. **A.** The structure of the α1 helix of EFR3A bound to TTC7B is shown on the top left. The AlphaFold models of TTC7B (middle) and TTC7A (right) are shown with all very low confidence regions (pLDDT values less than 50 being removed). AlphaFold predicts a capping helix in TTC7A at this binding interface, which has a reported phosphorylation site (Phosphosite) at S690 (colored yellow in the right panel). Additional reported TTC7A phosphorylation sites (Phosphosite) near this site are in predicted unstructured regions including S647, S678, S679, and T693. **B.** (L) EFR3A and (R) EFR3B phosphorylation sites (Phosphosite) mapped on the cryo-EM model of PI4KA complex-EFR3A C-terminus.

**Table S1.** Cryo-EM data collection, refinement, and validation statistics.

|  |  |
| --- | --- |
|  | PI4KA/TTC7B/FAM126A<br>(1-308)-EFR3A<br>C-terminus<br>EMD- 44413<br>PDB:9BAX |
| **Data collection and processing** |  |
| Magnification | 165,000 |
| Voltage (kV) | 300 |
| Electron exposure (e/ Å²) | 50 |
| Defocus range (µM) | 0.5-2.0 |
| Pixel size (Å) | 0.77 |
| Symmetry imposed | C1 |
| Initial particle images (no.) | 725,629 |
| Final particle images (no.) | 67,563 |
| Map resolution (Å) | 3.63 |
| FSC threshold | 0.143 |
| Map resolution range (Å) | 3.3-10 |
| **Refinement** |  |
| Initial model used (PDB) | 6BQ1/AlphaFold3 |
| Model Resolution (Å) | 3.48 |
| FSC threshold | 0.143 |
| Map sharpening B factor |  |
| Model composition |  |
| Non-hydrogen atoms | 40637 |
| Protein residues | 5082 |
| Ligands | 0 |
| *B*-factors |  |
| Protein | 242 |
| Validation |  |
| Mol probability score | 1.97 |
| Clashscore | 8.65 |
| Poor rotamers (%) | 1.48 |
| Ramachandran |  |
| Favoured | 94.50 |
| Allowed | 5.48 |
| Outliers | 0.02 |
| R.M.S. deviations |  |
| Bond lengths (Å) | 0.004 |
| Bond angles (°) | 0.565 |

**Table S2.**
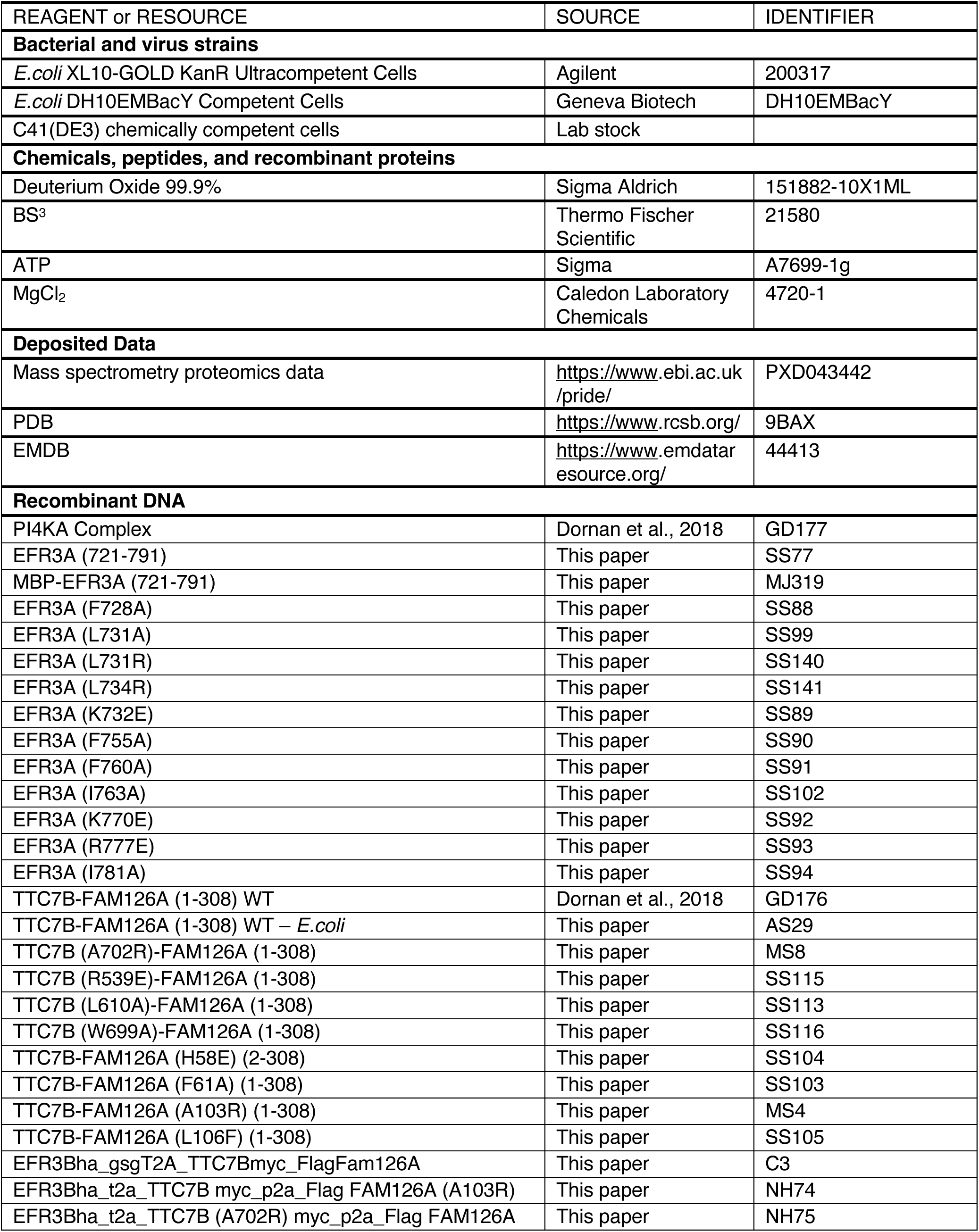
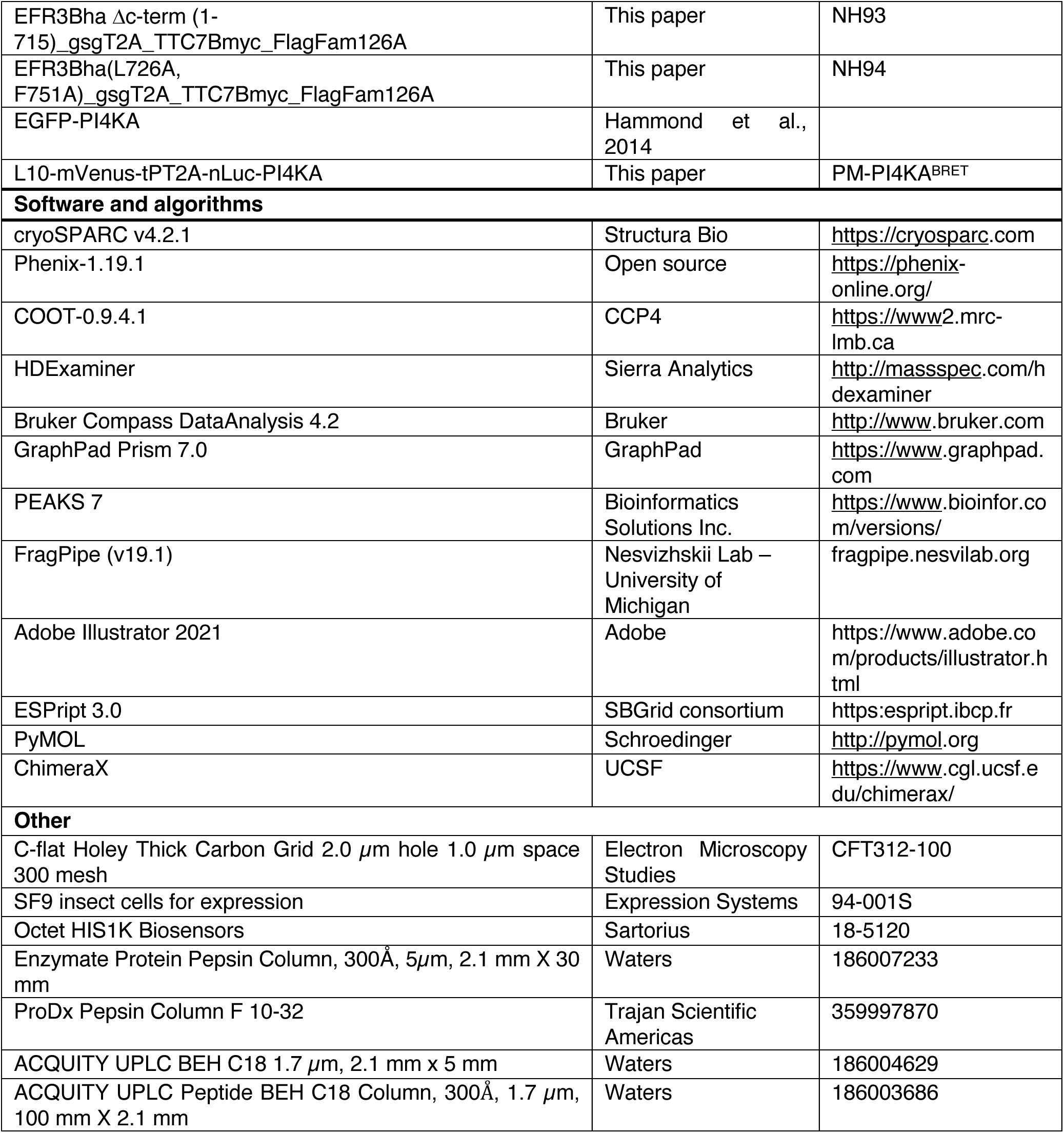
Key Reagent/Resources.

## Notes

### Summary of Updates

Minor alterations to the discussion to expand on isoform specific differences, and improvements to the computational modelling (alphafold3)

